# Bridging Morphology and Genomics: A rapid image-based assessment of genomic admixture in the endangered gayal (*Bos frontalis*)

**DOI:** 10.64898/2026.08.25.746947

**Authors:** Juntao Ma, Yuming Chen, Zhifan Guo, Jian Xiao, Jifan Yan, Haihong Wu, Jing Luo, Ya-Ping Zhang, Yan Li

## Abstract

The gayal (*Bos frontalis*) is an endangered semi-domesticated bovine species renowned for its high-quality beef. However, its semi-feral lifestyle, ongoing habitat fragmentation, and extensive genetic introgression from sympatric local cattle have led to dramatic population decline and severe erosion of purebred genetic integrity, posing substantial challenges to its conservation and utilization. To address the urgent demand for rapid, non-invasive, and field-compatible germplasm identification, we developed an integrated artificial intelligence (AI) framework that predicts genomic admixture composition from external morphological images. We constructed a comprehensive dataset comprising 6,245 morphological images and matched genomic sequences from 52 gayals maintained at the Yunnan Provincial Gayal Conservation Farms. Following a preliminary evaluation of nine deep learning models, five were incorporated into a anatomical segment-based multi-modal pipeline, among which Inception_V3 delivered the optimal overall performance. To enhance simultaneous extraction of local fine-grained features and global structural information, we further designed an innovative HybridInceptionViT model by integrating the multi-scale Inception module with the Vision Transformer (ViT) framework. This hybrid model significantly outperformed the baseline Inception_V3, boosting the accuracy of phenotype-derived prediction against genomic admixture estimate from 69.69% to 87.87% (absolute error <15%). This study establishes a practical, low-cost "phenotype-to-genotype" tool for rapid on-site gayal germplasm screening, offering a scalable strategy for the conservation and breeding management of endangered livestock, and holds broad application prospects for agricultural and livestock production systems.

**Highlights:** An integrative AI framework coupling image recognition with population genomic admixture was developed for rapid, non-invasive germplasm evaluation of the endangered gayal (*Bos frontalis*).

The novel HybridInceptionViT model synchronously captures local and global visual features, improving phenotype-based prediction accuracy of genomic ancestry from 69.69% to 87.87%, with absolute prediction error <15%.

This phenotype-to-genotype strategy provides a low-cost, field-adaptable tool for large-scale germplasm screening, supporting the conservation and breeding management of endangered livestock genetic resources.

## 1. Introduction

Gayal (*Bos frontalis*), also known as Mithun or Drung cattle, is an endangered semi-domesticated bovine species endemic to the rugged Eastern Himalayan foothills. Its distribution spans Yunnan and Xizang provinces of China, as well as India, Bangladesh, Myanmar, Nepal, and Bhutan (Dorji *et al*. 2021; Li *et al*. 2023; Simoons 1984). Adapted to rugged montane environments, gayal has evolved distinctive muscular characteristics, including small-diameter fibers with a fast-switch bias, fine and tender texture, low intramuscular fat content, mild flavor, and well-developed musculature (Ge *et al*. 1996; Han *et al*. 2025; Yunbo *et al*. 1998; Zhi *et al*. 2025; Li *et al*. 2023). These traits position it as a valuable genetic resource for improving beef quality and supporting the mountain husbandry economy.

However, due to its semi-wild lifestyle, insufficient conservation interventions, and anthropogenic disturbance, gayal populations have declined sharply (Chen *et al*. 2022; Uzzaman *et al*. 2014). More critically, extensive genetic introgression from local cattle has severely eroded the genetic integrity of purebred gayal (Li *et al*. 2023; Mei *et al*. 2016; Wu *et al*. 2018). Such genetic contamination has caused unstable phenotypic expression, particularly in economically important traits such as growth rate (Mukherjee *et al*. 2020; Uzzaman *et al*. 2014) and meat quality (Huang *et al*. 2025; Fan *et al*. 2005), thereby substantially threatening the conservation of authentic germplasm resources. Consequently, gayal has been listed as “endangered” in the World Watch List for Domestic Animal Diversity (FAO 2000), highlighting the urgency of targeted genetic conservation (Uzzaman *et al*. 2014). These challenges underscore the critical need for accurate germplasm identification to support the effective conservation and management of gayal genetic resources.

Conventionally, individual germplasm purity and ancestry composition are classically assessed using model-based clustering methods that assign individuals to *K* assumed ancestral populations (Pritchard *et al*. 2000). A series of computational tools have been developed for ancestry inference, including STRUCTURE (Pritchard *et al*. 2000), FRAPPE (Tang *et al*. 2005), ADMIXTURE (Alexander *et al*. 2009), FASTMIXTURE (Santander *et al*. 2024), etc.. Among these, ADMIXTURE is widely used to infer individual ancestry proportion using genome-wide single nucleotide polymorphism (SNP) data (Alexander *et al*. 2009). However, these approaches rely on tissue sampling and genomic sequencing, which require specialized expertise and may cause stress or injury to animals, particularly semi-feral gayal. Moreover, the relatively long turnaround time and high costs limit scalability, making them less suitable for rapid, large-scale germplasm assessment in practical farming systems. Therefore, the development of a rapid, accurate, and field-deployable identification method is urgently needed to support gayal germplasm conservation and targeted breeding.

The rapid development of CNN-based image recognition (Lecun *et al*. 1998; Krizhevsky *et al*. 2012) provides a promising technical foundation for such a rapid identification system. Current applications of image-based animal recognition primarily fall into two categories: species identification and individual recognition (Schneider *et al*. 2019).

In species identification, deep learning models have achieved stable and efficient performance, supporting automated wildlife monitoring and conservation. Early studies established automated pipelines for wildlife detection and classification using various CNN architectures (Nguyen *et al*. 2017). Subsequent comparisons between frameworks such as Faster R-CNN and YOLOv2 revealed trade-offs between detection accuracy and processing speed (Schneider *et al*. 2018). Building on this, de<u>ep</u> active learning improved recognition accuracy to ∼90% in complex natural environments (Norouzzadeh *et al*. 2021). Furthermore, the CNN-based ResNet-18 model attained 98% top-1 accuracy in automatic animal identification from camera-trap images in the United States (Tabak *et al*. 2019). These studies demonstrate that deep learning-enabled image recognition has become a robust tool for ecological monitoring and wildlife research (Oliveira *et al*. 2021).

Compared with species identification, individual recognition is more challenging due to high intraspecific morphological similarity. Nevertheless, distinctive visual patterns, such as zebra stripes, cheetah spots, or fish fin markings enable reliable individual discrimination (Bolger *et al*. 2012). Early studies applied traditional machine learning methods, including computer vision techniques and support vector machines, to identify individual Holstein cattle based on coat patterns (Li *et al*. 2017). Recent deep learning approaches have substantially improved the accuracy and scalability of individual recognition. For instance, VGGNet achieved 95% accuracy in individual giant panda identification using 65,000 facial images (Hou *et al*. 2020). The same model attained 98.7% accuracy in beef cattle individual identification using mouth patterns (Li *et al*. 2022). Using EfficientNetV2, Takaya et al. further pushed the boundary by achieving 99.86% accuracy in identifying endangered Japanese giant salamanders from skin patterns (Takaya *et al*. 2023). These studies highlight the value of local features in individual recognition. Notably, optimal model selection depends on by task-specific factors, such as animal body size, background complexity, and targeted anatomical regions (Vidal *et al*. 2021). Accordingly, integrating different deep learning architectures is preferable to improve performance in specialized recognition tasks by enabling more comprehensive extraction of discriminative phenotypic traits (Xu *et al*. 2024). Despite the rapid progress in visual recognition, the use of image-based approaches to infer animal germplasm composition remains largely unexplored.

Against this background, integrating CNN-based phenotypic features with ADMIXTURE-derived individual genetic components makes it possible to predict germplasm composition directly from morphological images. We therefore developed a novel tool for evaluating gayal germplasm admixture using image-based morphological recognition. This method requires only multi-angle photographs and can complete phenotypic feature extraction and germplasm inference within minutes. It is non-invasive, convenient, and time-saving, making it ideal for large-scale field applications. Consequently, this approach provides a practical solution for on-site germplasm conservation and breeding guidance, with promising applications across various livestock production and conservation systems.

## 2. Materials and methods

### 2.1. Whole-genome sequencing, variant calling, and ADMIXTURE analysis

Ear tissues from multiple populations inhabiting Fenghuang Mountain, Lushui City, Yunnan Province, China were collected. Samples were immediately frozen in liquid nitrogen and stored at −80 °C until DNA extraction. All animal experimental procedures were approved by the Institutional Animal Care and Use Committee of Yunnan University (Approval No. YNU20251450). Whole-genome sequencing of 67 gayal was performed at Novogene Bioinformatics Technology Co., Ltd using the Illumina platform with a 150 bp paired-end (PE150) strategy. Combined with whole-genome resequencing data of additional 47 cattle downloaded from the NCBI Sequence Read Archive (SRA)(**Appendix A and B)**, a total of 114 individuals were retrieved for subsequent genomic analysis.

Raw reads were quality-filtered using Trimmomatic v0.39 (Bolger *et al*. 2014) to remove adapter sequences and low-quality bases. Reads with average quality scores below threshold or shorter than 50 bp were discarded. Clean reads were processed using the BaseNumber germline variant detection pipeline, following the Genome Analysis Toolkit (GATK) (4.1.2.0) (Van der Auwera *et al*. 2020) Best Practices for single nucleotide polymorphism (SNP) detection, including read alignment, duplicate marking, and variant calling. Reads were aligned to the gayal reference genome (GCA_043643345.1) using BWA 0.7.17 (Li *et al*. 2009). PCR duplicates were removed and GATK HaplotypeCaller was used to generate gVCF files. Joint genotyping was then performed to obtain the final VCF dataset. SNPs were filtered using GATK VariantFiltration with the following criteria: QD < 2.0, QUAL < 30.0, SOR > 3.0, FS > 60.0, MQ < 40.0, MQRankSum < −12.5, and ReadPosRankSum < −8.0.

Further filtering was performed using PLINK 1.90 (Purcell *et al*. 2007) to remove SNPs with low minor allele frequency (MAF < 0.05), high missing rate (geno > 0.1), deviation from Hardy-Weinberg equilibrium (HWE P < 0.0001), or strong linkage disequilibrium (--indep-pairwise 50 10 0.1). Sex chromosomes were excluded from downstream analyses. Population genetic structure was inferred using ADMIXTURE 1.3.0 (Alexander *et al*. 2009) based on 114 individuals **(Appendix C)**. The optimal ancestral population number *K* was determined by cross-validation, and ancestry coefficients (Q matrix) were visualized using TBtools-Ⅱ2.3.3.0 (Chen *et al*. 2023).

### 2.2. Image collection and preprocessing

Gayal individuals with matched resequencing data were photographed using the standard photo mode of a Xiaomi 14 smartphone. Two rounds of image collection were conducted. The first dataset comprising 715 images from 15 gayal individuals **(Appendix D)** was used for model pre-screening. Based on preliminary model training and evaluation, a second, more standardized dataset consisting of 6,245 images from 52 gayal individuals **(Appendix D)** was collected following strict criteria: (1) relatively balanced image counts among individuals; (2) coverage of eight viewing orientations (front, right front, left front, right side, left side, back, right back, and left back); (3) no obvious obstructions in front of the individual; and (4) minimal co-occurrence of multiple individuals in single frames. After removing low-quality images and those of juvenile individuals, photographs from 41 gayal individuals were retained and randomly partitioned into a training set (33 individuals) and a testing set (8 individuals) at a ratio of 4:1.

To facilitate targeted local feature extraction, all images from the second dataset were manually cropped into three anatomical regions: head, body, and limbs. The anatomical region-based cropped images were also divided into training and validation sets at a 4:1 ratio. Automatic object detection algorithms such as YOLO-based methods (Redmon *et al*. 2016)) were not used for image cropping, mainly for the following reasons: (1) object detection algorithms require extensive time-consuming manual annotation; (2) such algorithms are incompatible with the local feature-based multi-modal pipeline adopted in this study; (3) bounding boxes generated by object detection algorithms may introduce background noise. Accordingly, manual cropping was performed using maximal rectangular bounding boxes to retain complete morphological features while excluding irrelevant background.

### 2.3. Model pre-screening

To identify suitable architectures for gayal germplasm evaluation, nine mainstream deep learning models were pre-screened using the first image dataset, including ResNet (ResNet50, ResNet101, ResNet152)(He *et al*. 2016), Vision Transformer (ViT)(ViT_b_16, ViT_b_32)(Dosovitskiy *et al*. 2020), VGG16 (Simonyan *et al*. 2014), EfficientNet (EfficientNet_b7, EfficientNet_v2_l)(Tan *et al*. 2019, 2021), and Inception_V3 (Szegedy *et al*. 2016). During preprocessing, the short side of each image was resized to 512 pixels while maintaining aspect ratio. Data augmentation was applied to improve model robustness, including random 448×SimSun>448 cropping for feature extraction and horizontal flipping with 50% probability to increase sample diversity. Images were then converted to tensors and normalized using statistics from the ImageNet (Deng *et al*. 2009) pre-training dataset to support transfer learning.

Model training for gayal image classification was performed using cross-entropy loss (Goodfellow *et al*. 2016), the standard objective function for multi-classification. Cross-entropy loss quantifies the discrepancy between predicted and true label distributions; unlike mean square error (MSE)(Goodfellow *et al*. 2016), it updates gradient magnitude in proportion to prediction confidence and imposes stronger penalties for misclassifications, thereby accelerating convergence and enhancing inter-class discriminative feature learning.

Parameters were optimized using stochastic gradient descent (SGD) with momentum (Goodfellow *et al*. 2016). The initial learning rate was set to 0.002 and decreased to 0.001, 0.0005, and 0.0001 at epochs 20, 30, and 50, respectively. A weight decay coefficient of 5×10⁻⁵ was applied to mitigate overfitting. SGD with momentum was adopted for its low computational cost and stable convergence performance, while weight decay improved generalization by reducing model complexity.

### 2.4. Multi-modal training pipeline

Following identification of the optimal baseline model, a multi-modal training pipeline was implemented using anatomically segmented images from the second dataset. Two training strategies were compared: transfer learning and training from scratch. Transfer learning leverages feature representations learned from large-scale general datasets to improve performance on targeted tasks, whereas training from scratch allows the model to learn task-specific features directly from the target dataset. Comparative evaluation of these strategies enabled us to determine the most effective paradigm for gayal germplasm identification. All pre-trained feature representations were obtained from the official PyTorch repository (Paszke *et al*. 2019), originally trained on the ImageNet dataset to provide general visual feature priors.

Training of the multi-modal pipeline followed consistent hyperparameters as described above. Cross-entropy loss was used as the objective function, and SGD with momentum was used for optimization. Transfer learning was implemented by initializing pre-trained ImageNet weights and fine-tuning on the gayal dataset. Models were trained on anatomical region-based cropped images of 33 individuals (randomly selected from the second dataset) with matched admixture values (see Section 2.1) for a maximum of 80 epochs, with early stopping applied when both training and validation accuracy stabilized to prevent overfitting.

During model inference, a fixed random seed was set to ensure experimental reproducibility. Both scratch-trained and transfer learning-trained models were loaded from locally saved checkpoints. The final fully connected layer of each model was modified to match the number of gayal identification classes. Models were switched to evaluation mode during testing to ensure stable prediction. Individual identity was predicted as the class with the highest posterior probability (top-1 class).

To evaluate model generalization and reduce potential overfitting, a sampling-based validation strategy was applied to all trained region-based models. For each model, three images were randomly selected from three distinct individuals, and predicted labels were compared against ground truth. This sampling inspection was performed independently for each trained architecture.

### 2.5. Model attention visualization

Deep learning models are often regarded as black boxes. To verify whether models focused on biologically meaningful morphological features rather than background noise, model interpretability was analyzed using Gradient-weighted Class Activation Mapping (Grad-CAM) algorithm (Selvaraju *et al*. 2017). Grad-CAM generates a visual heatmap by computing channel-wise importance weights from class-specific gradients, producing weighted sums of convolutional feature maps while suppressing negative contributions. For visualization, three images from three individuals were randomly selected for each anatomical region (head, body, and limbs). Attention maps were generated for both transfer-learned and scratch-trained models. Attention distributions were compared to evaluate differences in feature localization.

To ensure consistent interpretability across multiple architecture, the last convolutional layer of each models was used to maximize the receptive field and ensure full image coverage. Pytorch Hook functions (Paszke *et al*. 2019) were used to monitor module forward propagation and tensor gradient computation during visualization.

### 2.6. Construction and application of HybridInceptionViT

To further improve feature representation and identification accuracy for gayal germplasm evaluation, a novel hybrid deep learning architecture named HybridInceptionViT was developed by integrating the complementary advantages of InceptionV3 (Szegedy *et al*. 2016) and ViT (Dosovitskiy *et al*. 2020). The model combines the strong local and multi-scale feature extraction capability of Inception with the powerful global context modeling of ViT, enabling joint learning of fine-grained local features and holistic global structure representations.

The architecture consists of three modules: an Inception feature extractor, a ViT feature extractor, and a feature fusion component. In the Inception branch, the original fully connected layer of InceptionV3 was replaced with an identity mapping to output a 2048-dimensional feature vector. High-frequency convolution operations were introduced into key layers (Mixed_5b, Mixed_5c, and Mixed_5d) to enhance fine-scale feature extraction. In parallel, the ViT branch processes image patches through a Transformer encoder to generate a 768-dimensional global feature representation.

Features from the two branches were concatenated into a 2816-dimensional combined vector, which was subsequently projected to the target number of classes through two fully connected layers equipped with Batch Normalization, ReLU activation, and Dropout regularization to improve generalization. Model training, image preprocessing, and training strategies (transfer learning vs. scratch) remained consistent with previous settings. The trained HybridInceptionViT model was applied for prediction by selecting the class with the highest predicted probability.

To interpret the feature learning mechanism of the hybrid model, Grad-CAM was applied to visualize spatial attention across different modules. Heatmaps were generated from the final convolutional layer of the Inception branch, the ViT feature layer, and the fused feature layer to evaluate the respective contributions of local and global features to final prediction.

### 2.7. Multi-modal pipeline prediction and fitting with genomic data

To evaluate the consistency between image-based predictions and genomic germplasm composition, uncropped full-body images of 33 individuals from the training dataset and 8 individuals from the independent testing dataset were directly input into three anatomical region-based prediction pipelines (body, head, and limb models). Each pipeline generated an independent prediction score for its respective anatomical segment. For each gayal individual, five representative images were used for prediction, and the prediction with the highest confidence was retained as the final output. The predicted scores from the three pipelines (Body_Pre, Head_Pre, and Limb_Pre) were then regressed against the genomic ancestry germplasm composition estimated from whole-genome sequencing.

Linear regression, nonlinear regression, and machine learning-based fitting approaches were tested. Among them, a ternary multiple linear regression model achieved the best fitting performance. The model is formulated as follows:

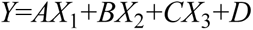

where *X*1, *X*2, and *X*3 represent prediction outputs of the limb, head, and body pipelines, respectively. Model performance was evaluated using the coefficient of determination (*R*²), adjusted *R*², *F*-statistics, and corresponding *P*-values, with multi-modal phenotypic predictions as independent variables and genomic admixture proportions as the dependent variable.

## 3. Results

### 3.1. Population genetic structure

Since population genetic admixture analysis provideds a reliable benchmark for germplasm identification, and this study aimed to build a robust phenotype-genotype mapping, we first performed population structure analysis for the gayal individuals included in the model training set. Genetic structure was inferred using ADMIXTURE with *K* values from 2 to 10. Cross-validation error was minimized at *K*=3 (results for *K*=4 were also presented in **Appendix E** for comparison), indicating that three ancestral populations optimally explained the observed genetic structure. At *K*=3, individuals clustered into three distinct groups corresponding to gayal, zebu, and taurine cattle **(Appendix E)**. Although gayal individuals formed a well-defined genetic cluster, varying levels of zebu ancestry were detected in several individuals, indicating prominent zebu introgression within the gayal population **(Appendix E)**.

### 3.2. Image processing and dataset construction

The initial dataset included images from 15 gayal individuals. Three individuals were excluded due to an insufficient number of available images, resulting in a final preliminary dataset of 685 images from 12 individuals **(Appendix F)**. These images exhibited substantial unbalanced representation among individuals **(Fig.1-A)** and frequently contained obstructions or multiple individuals within a single frame. Therefore, this dataset was exclusively used for preliminary screening and comparison of deep learning models for individual identification.

**Fig. 1.**
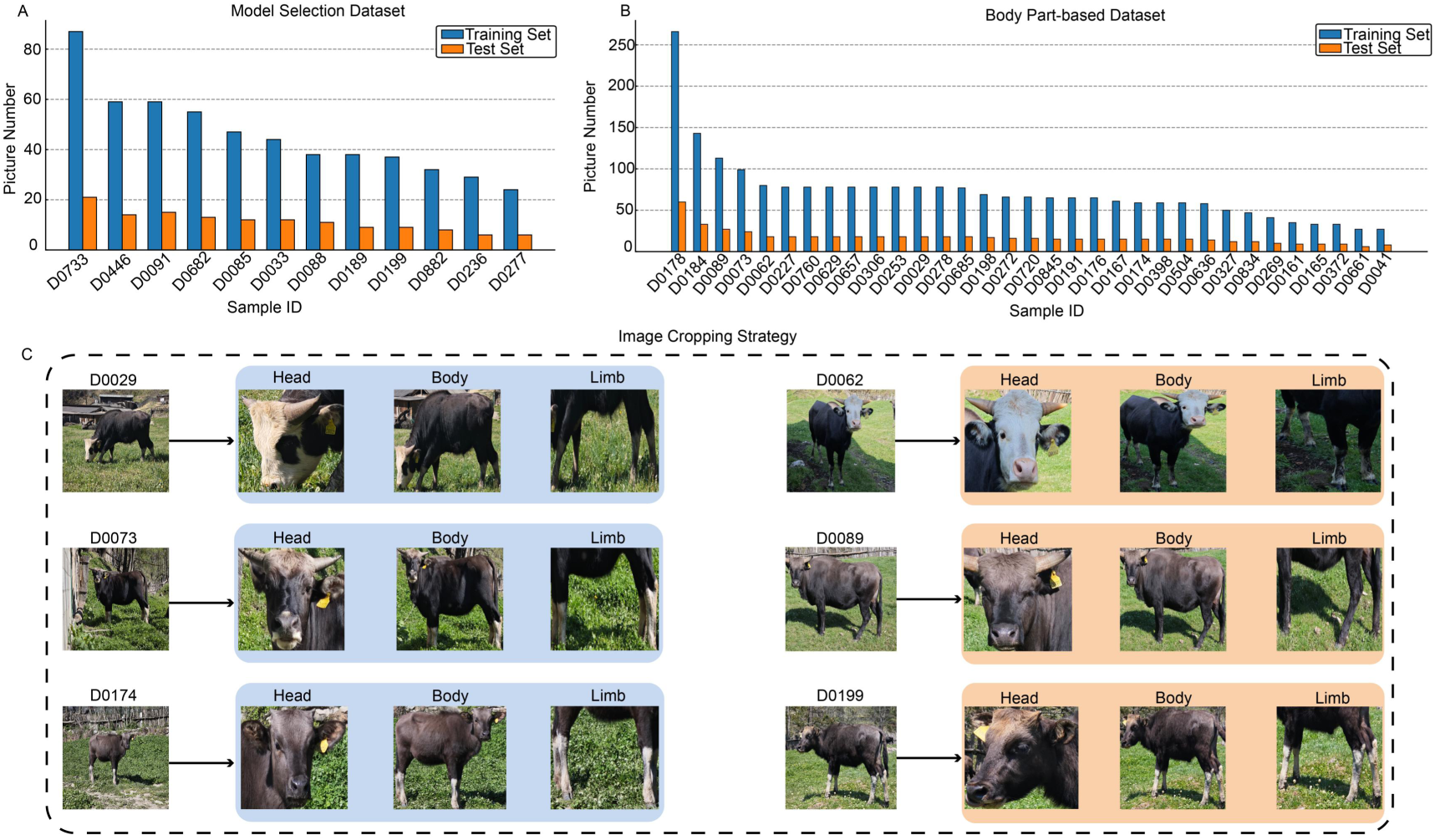
Construction of the gayal image dataset and part-based preprocessing strategy. A, Composition of the dataset used for model selection. Images from 12 gayal individuals were used, and the training and test sets were constructed at a 4:1 ratio based on images of each individual. B, Composition of the body part-based dataset used for model training. After screening, images from 33 individuals were included, and the dataset was split into training and test sets at a 4:1 ratio according to individual identity. C, Illustration of the image cropping strategy used to generate part-based datasets. Raw field images were manually cropped into three anatomical regions (head, body, and limb); six representative individuals are shown.

A second high-quality dataset was subsequently established under stricter image acquisition criteria, consisting of 6,245 images from 52 individuals **(Appendix D)**. After removing low-quality images and those of juvenile individuals, photographs from 41 gayal individuals were retained and randomly partitioned into a training set (33 individuals) and a testing set (8 individuals) at a ratio of 4:1. Within the training set, the anatomical region-based cropped images were also divided into training and validation sets at a 4:1 ratio **(Fig. 1-B; Appendix G-I)**. Although slight imbalance in image counts persisted among certain individuals (e.g., D0178 and D0041), the distribution between training and validation sets was strictly controlled **(Fig. 1-B)**. To facilitate local feature-based phenotypic extraction, images were manually cropped into three anatomical subsets representing head, body, and limbs **(Fig. 1-C; Appendix J)**. These curated datasets were subsequently used for multi-model training and evaluation.

### 3.3. Model Training

Following dataset construction, nine mainstream deep learning models were trained on the first 685 image dataset to identify the most appropriate architecture fo<u>r further</u> <u>multi-modal pipeline construction</u> **(Table 1)**. Among all tested architectures (EfficientNet_b7, EfficientNet_v2_l, ResNet50, ResNet101, ResNet152, Inception_V3, VGG16, ViT_b_16, ViT_b_32), the EfficientNet series achieved excellent performance but suffered from severe underfitting (**Appendix K**), making them unsuitable for our relatively small dataset. For the ResNet series, ResNet50 achieved the highest validation accuracy (98.5%), followed by ResNet101 (97.7%) and ResNet152 (97.0%)**(Table 1)**. ResNet networks are primarily modified by increasing convolutional depth rather than altering structural modules. Given the insignificant differences of layer count between ResNet_101 and ResNet_50, which may lead to analogous feature extraction patterns, we chose ResNet50 and ResNet_152 (representing the shallowest and deepest networks) as the candidates for downstream pipeline construction.

**Table 1.**
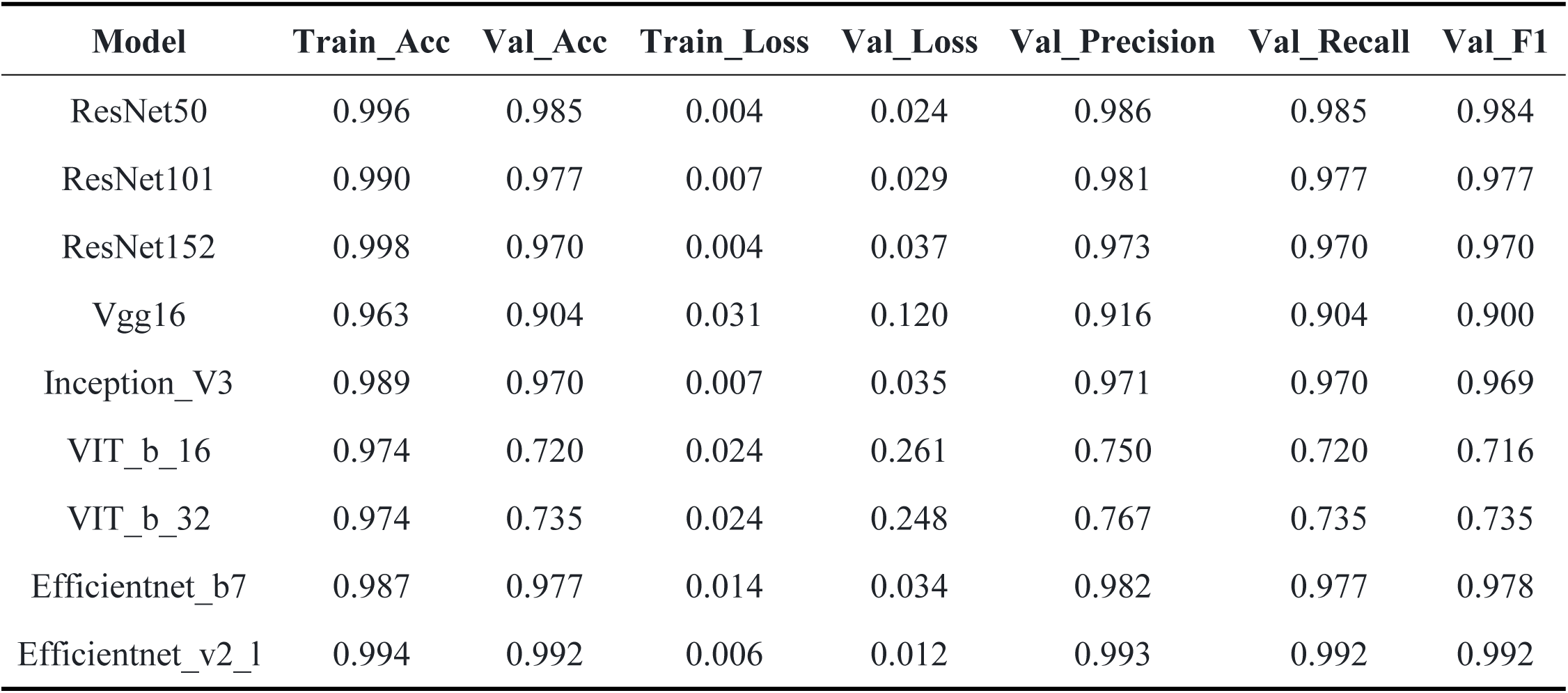
Training Performance of Nine Models on the Model Selection Training Set.

For other architectures, Inception_V3 exhibited stable and favorable performance despite its relatively small parameter size, owing to its distinctive multi-branch feature extraction design. Although VGG16 did not performed optimally, its large parameter size might support complex feature learning in expanded dataset. Accordingly, we also retained Inception_V3 and VGG16 as candidates.

Vision Transformer (ViT) models employ a powerful self-attention mechanism, which represented a distinct paradigm for feature extraction in image recognition. However, ViT series performed poorest in the pre-screening **(Fig. 2-C)**, probably due to the small size in the first dataset that was far from robust ViT training (Touvron *et al*. 2021b). Nevertheless, the self-attention mechanism remains highly valuable; we therefore integrated ViT into a hybrid model for improved performance on the larger second dataset. Collectively, five models, including ResNet50, ResNet152, VGG16, Inception_V3, and ViT_b_32, were chosen for further pipeline construction.

**Fig. 2.**
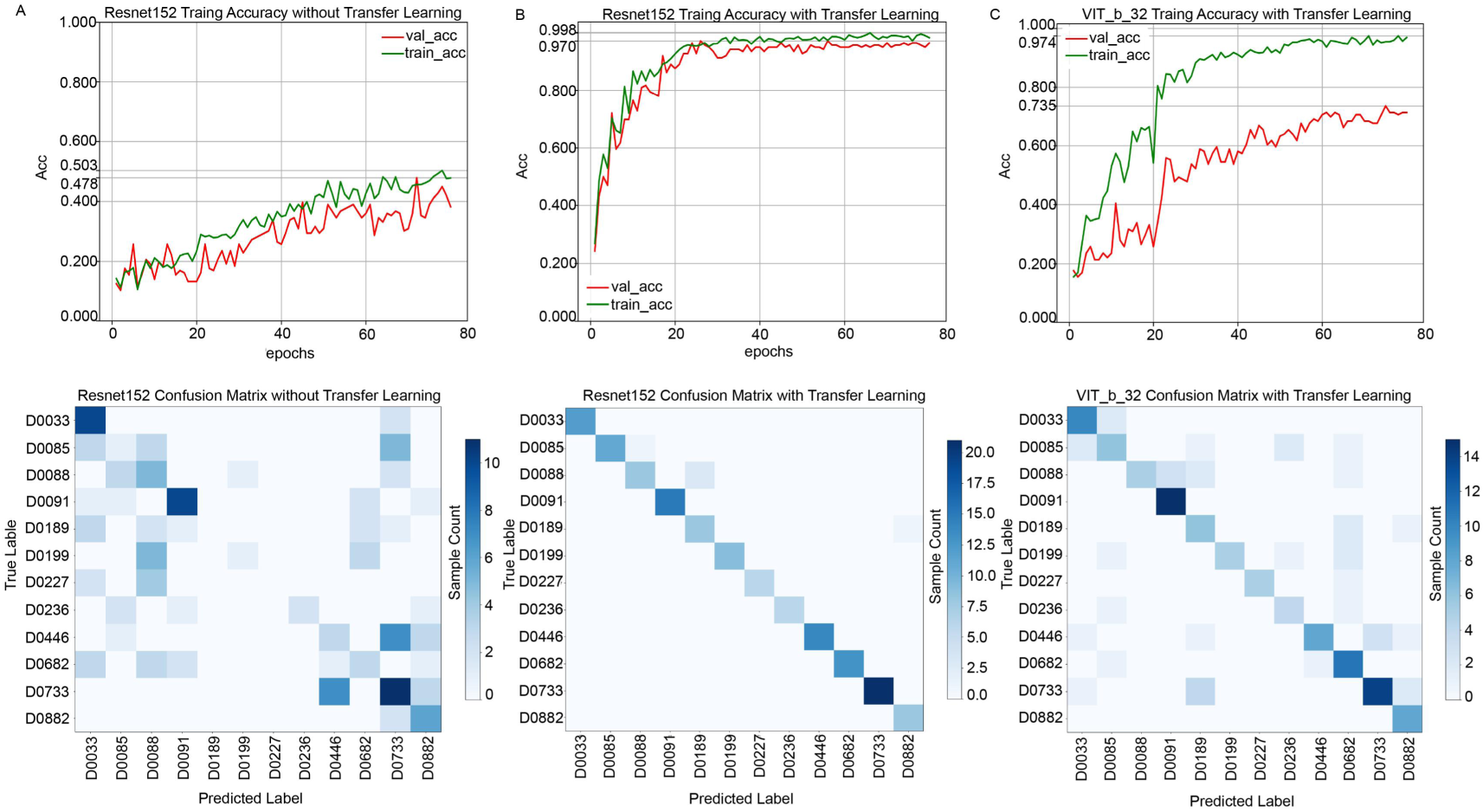
Performance evaluation of representative deep learning models for Gayal identification. A, Performance of the ResNet152 (no transfer learning) on the model selection dataset: training/validation accuracy in the upper panel, corresponding confusion matrix in the lower panel. B, Performance of the ResNet152 model with transfer learning. C, Performance of the ViT_b_32 with transfer learning.

The multi-model pipeline was trained on approximately 9000 cropped anatomic region-based images from 33 individuals in the second dataset. Two training strategies were compared: transfer learning and training from scratch. Overfitting as a source of inflated accuracy was ruled out through independent prediction validation and Grad-CAM attention heatmap analysis. In all experiments, models trained with transfer learning consistently outperformed those trained from scratch **(Table 2)**. The relatively poor classification performance of scratch-based models was likely due to limited dataset size. For example, scratch-based ResNet152 model failed to achieve effective individual discrimination, as reflected in its confusion matrix **(Fig. 2-A)**. In contrast, transfer learning-based models **(Fig. 2-B)** substantially improved the classification accuracy and produced reliable predictive performance evidenced by the confusion matrix. With the exception of ViT, transfer learning improved training accuracy by nearly 10% under identical settings and substantially reduced the performance gap between training and validation sets.

**Table 2.** Head data Training Results of Scratch Training vs. Transfer Learning for 5 Models (TL = Transfer Learning)

| Model | Train_Acc | Val_Acc | Train_Loss | Val_Loss | Val_Precision | Val_Recall | Val_F1 |
| --- | --- | --- | --- | --- | --- | --- | --- |
| ResNet50<sup>s</sup> | 0.878 | 0.930 | 0.337 | 0.289 | 0.937 | 0.930 | 0.928 |
| ResNet50_TL | 0.995 | 0.998 | 0.003 | 0.003 | 0.998 | 0.998 | 0.998 |
| ResNet152 | 0.853 | 0.900 | 0.312 | 0.305 | 0.933 | 0.900 | 0.901 |
| ResNet152_TL | 0.997 | 0.998 | 0.002 | 0.002 | 0.998 | 0.998 | 0.998 |
| VGG16 | 0.896 | 0.952 | 0.086 | 0.042 | 0.956 | 0.952 | 0.950 |
| VGG16_TL | 0.982 | 0.986 | 0.015 | 0.008 | 0.987 | 0.986 | 0.986 |
| Inception_V3 | 0.871 | 0.932 | 0.115 | 0.060 | 0.938 | 0.932 | 0.931 |
| Inception_V3_TL | 0.990 | 0.994 | 0.010 | 0.003 | 0.995 | 0.994 | 0.994 |
| VIT_b_32 | 0.972 | 0.857 | 0.021 | 0.119 | 0.868 | 0.857 | 0.856 |
| VIT_b_32_TL | 0.994 | 0.945 | 0.005 | 0.046 | 0.947 | 0.945 | 0.944 |

### 3.4. Model validation and feature interpretability

All five selected architectures achieved high classification performance. Among them, ResNet_152 fine-tuned via transfer learning demonstrated the best overall performance, while Inception_V3 exhibited stable performance under both training strategies. Training dynamics revealed clear strategy-dependent differences. Training curves showed that scratch-trained models converged more slowly and displayed more volatile loss trajectories compared with transfer learning models **(Figs. 3-A–B)**. In particular, scratch-trained InceptionV3 exhibited a steep learning curve and greater loss fluctuation despite eventually reaching acceptable accuracy **(Figs. 3-A)**. Within the 80-epoch training schedule, transfer learning enabled faster convergence and more stable optimization, whereas scratch training required greater computational time to achieve comparable performance **(Figs. 3-B)**.

**Fig. 3.**
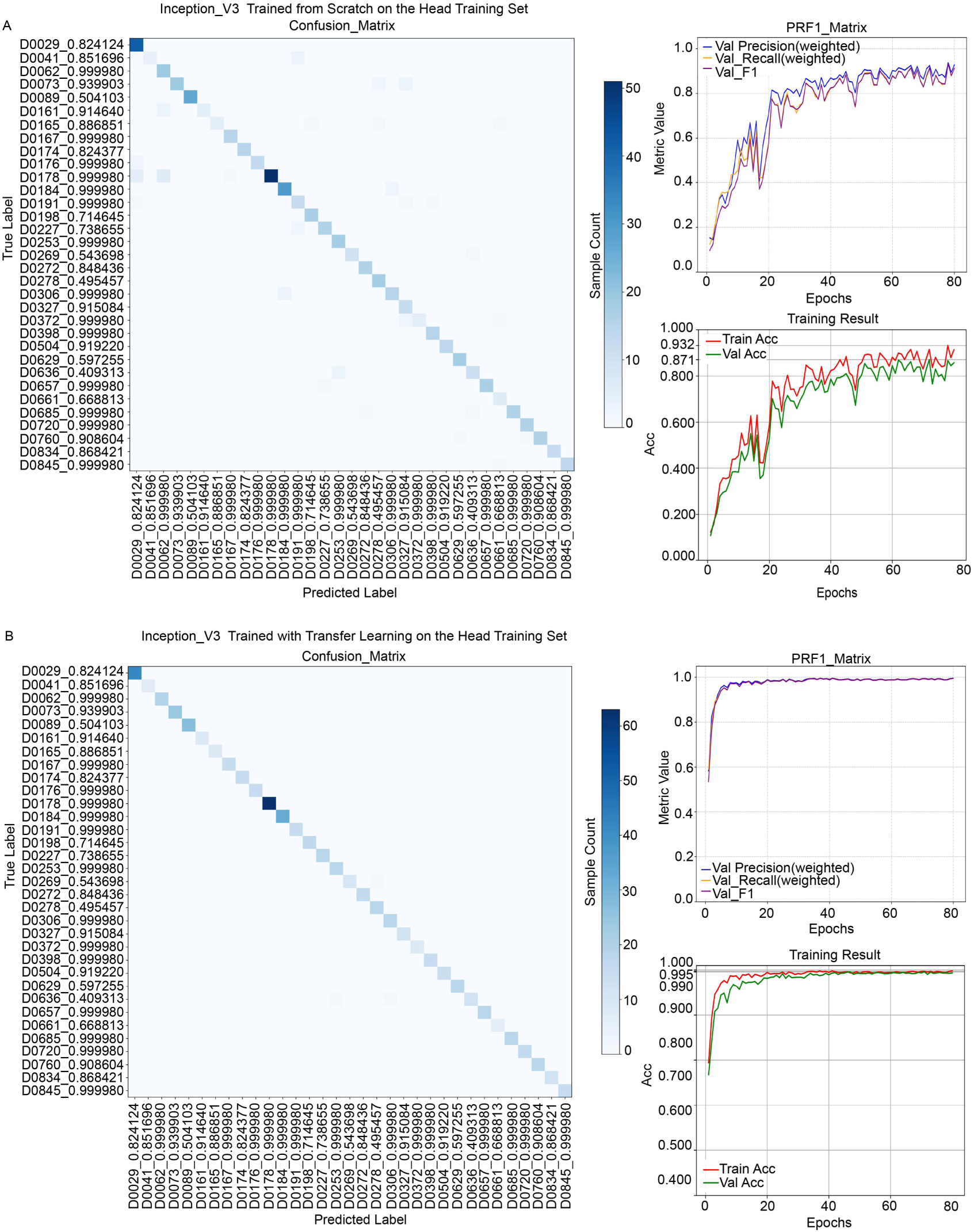
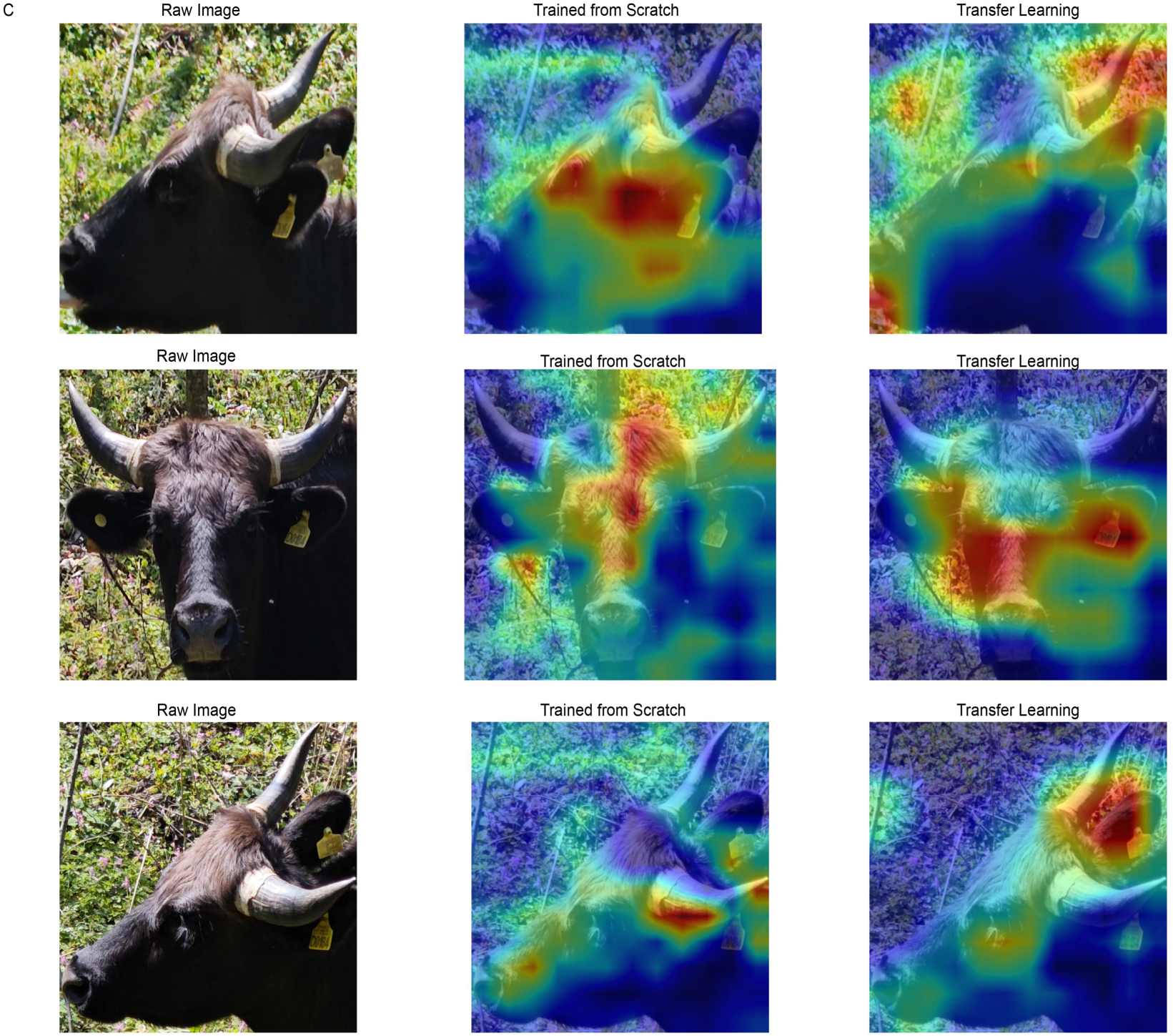
Performance comparison and attention visualization of InceptionV3 models trained with different strategies on the head-region dataset. A, Training metrics of Inception_V3 with scratch training on the head region training set. B, Training metrics of Inception_V3 with transfer learning on the head region training set. C, Grad-CAM visualization for three head images of individual D0184 using Inception_V3 based on the two training strategies.

Sampling-based inspection revealed distinct differences in model generalization. Among all trained models, Inception_V3 achieved 100% accuracy across all sampled test instances. Based on these results, Inception_V3 was selected as the base architecture for the recognition pipeline with three anatomic regions.

Grad-CAM visualization uncovered substantial differences in spatial attention between training strategies. Transfer learning models consistently focused on biologically meaningful anatomical regions of gayal, whereas scratch-trained models allocated considerable attention to background areas. Visualization results for three body parts of individual D0029 **(Appendix L)** showed that the transfer learning model more accurately concentrated on targeted body regions. Particularly, the scratch model distributed attention randomly across the head region, whereas the transfer learning model focused more precisely on phenotypic features, consistent with dataset design intent. Visualization of the head region for individual D0184 was even more distinct **(Fig. 3-C)**: The transfer learning model showed highly localized activation on the head with minimal background interference.

The limb dataset presented the greatest challenge due to the relatively small size of legs within full-body images. As expected, attention maps indicated partial interference from background noise **(Appendix M)**. Nevertheless, the transfer learning model accurately localized attention to the legs without obvious background activation, whereas scratch-trained models displayed dispersed attention patterns **(Appendix M)**. Overall, attention visualization across all three anatomical datasets supported the adoption of transfer learning for pipeline construction.

### 3.5. HybridInceptionViT model training and validation

To enhance feature extraction capability of the InceptionV3-based pipeline, we innovated a hybrid model, HybridInceptionViT, which integrated the self-attention mechanism of ViT **(Fig. 6-B)**. Comparison between training results in **Table 3** and baseline performances in **Table 2** demonstrated that HybridInceptionViT model achieved consistently high accuracy under both transfer learning and scratch training. This performance indicated that its combined feature extraction strategy is better suited to our dataset. As shown in **Fig. 4-A** **and B**, the hybrid architecture demonstrated faster convergence and more stable training dynamics than conventional single models. In particular, even when trained from scratch, the hybrid model reached stability within fewer epochs, indicating improved training efficiency.

**Fig. 4.**
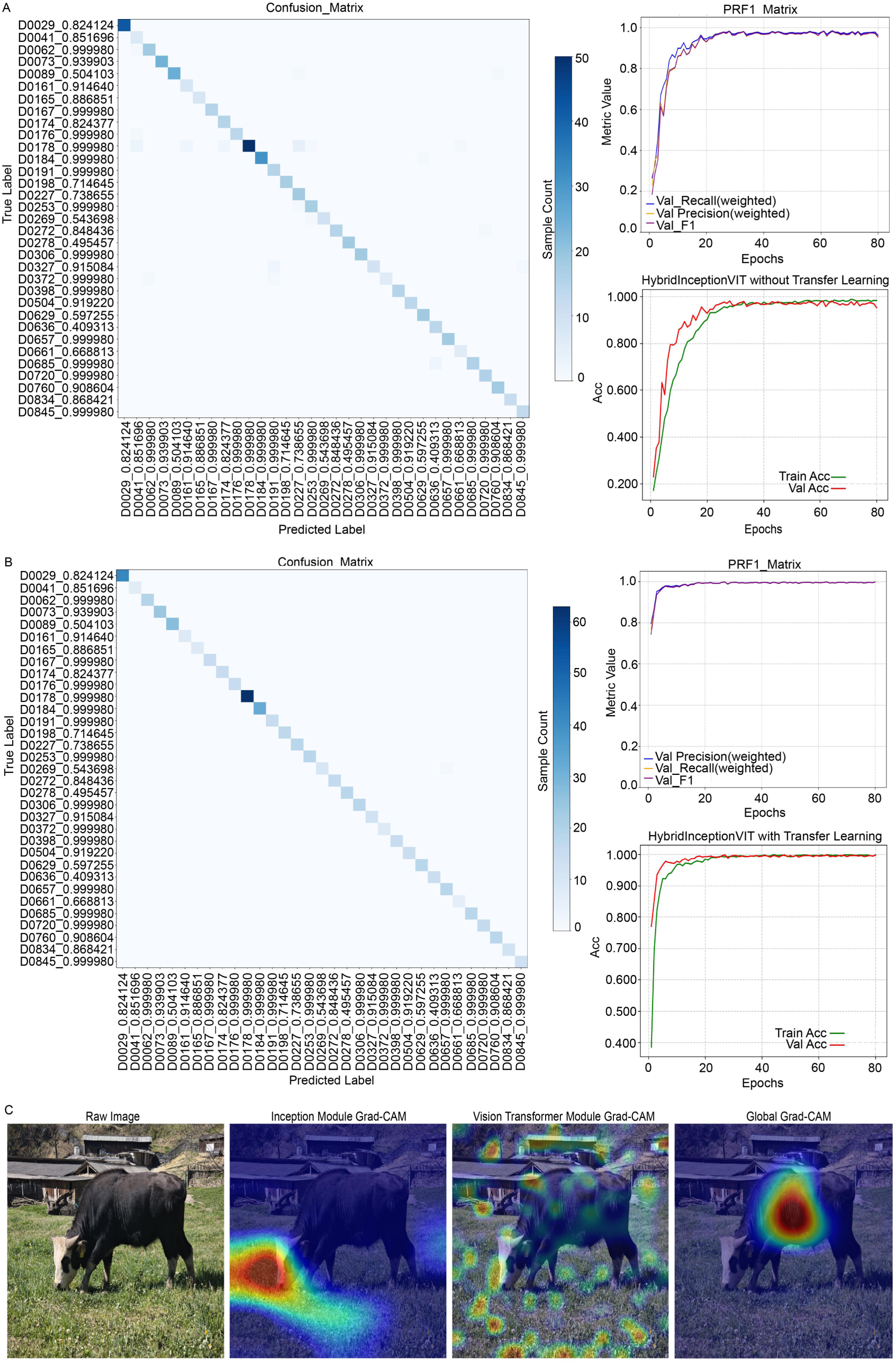
Training performance and attention visualization of the HybridInceptionViT model on the part-based dataset. A, Training performance of the HybridInceptionViT with scratch training on the head region training set. B, Training performance of the HybridInceptionViT with transfer learning on the head region training set. C, Comparative Grad-CAM visualization of the Inception branch, ViT branch, and feature fusion layer for a single raw image of individual D0029 using HybridInceptionViT with scratch training on the head region training set.

**Fig. 6.**
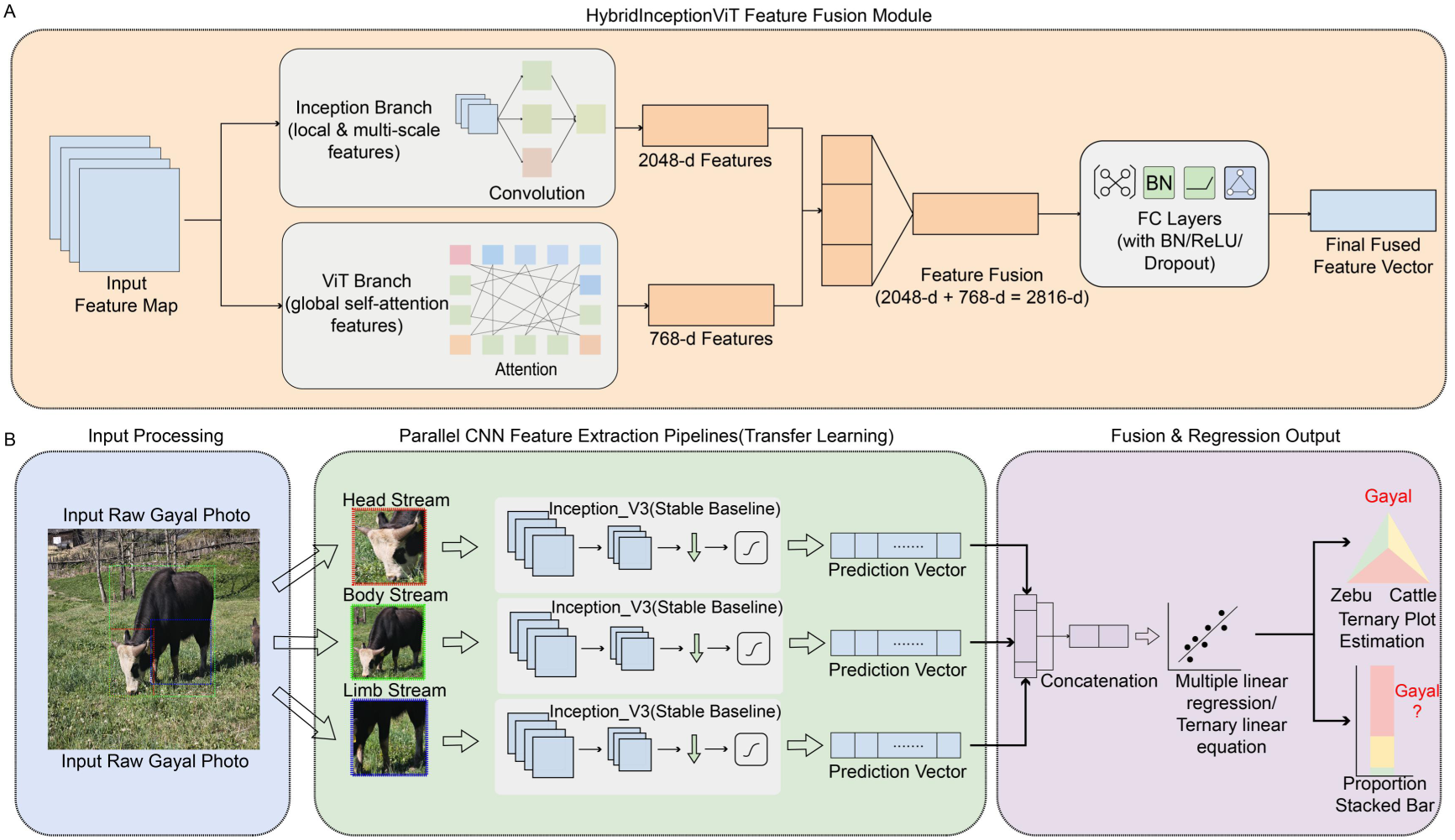
Deep-learning framework for image-based Gayal germplasm identification. A, Architecture of the HybridInceptionViT feature-fusion model. The model combines an Inception branch for local and multi-scale feature extraction with a ViT branch for global context modeling, and the concatenated features are processed by fully connected layers to generate the final prediction. B, Schematic diagram of germplasm fitting from gayal images. Raw images were partitioned into three anatomical regions (head, body, and limb), processed in parallel using transfer-learning models based on InceptionV3, and the resulting prediction vectors were concatenated and integrated to estimate the final germplasm composition.

**Table 3.** HybridInceptionViT Training Results on Head Data of Scratch Training vs. Transfer Learning (TL = Transfer Learning)

| HybridInception<br>ViT | Train_Acc | Val_Acc | Train_Loss | Val_Loss | Val_Precision | Val_Recall | Val_F1 |
| --- | --- | --- | --- | --- | --- | --- | --- |
| Without_TL | 0.988 | 0.981 | 0.048 | 0.070 | 0.983 | 0.981 | 0.981 |
| TransferLearning | 0.999 | 0.998 | 0.005 | 0.013 | 0.998 | 0.998 | 0.998 |

Grad-CAM visualization revealed distinct attention distribution between model components **(Fig. 4-C)**. The Inception branch primarily focused on local, fine-grained regions of the image, while the ViT branch distributed attention more broadly across the entire image **(Fig. 4-C)**. Although the ViT branch exhibited mild background dispersion, it effectively covered the entire body of the gayal individual, consistent with the original design. Following feature fusion, model attention became more tightly concentrated on the gayal with reduced background interference.

### 3.6. Fitting of model predictions and genomic data

To evaluate whether image-based predictions could reliably reflect genomic germplasm composition, prediction outputs from the three anatomical pipelines were fitted to genomic admixture proportions using a ternary linear regression model **(Fig. 6-B)**. Tables 4 and 5 summarized the fitting performance between the genomic estimates and phenotype-based predictions from the Inception-V3 and HybridInceptionViT models, respectively.

**Table 4.** Authentic Germplasm Composition and Part-specific Germplasm Prediction Results with Inception_V3 Pipeline.

| True<br>value | LIMB_Pre | HEAD_Pre | TOTAL_Pre | Predict<br>Value | Deviation |
| --- | --- | --- | --- | --- | --- |
| 0.824124 | 0.409313 | 0.99998 | 0.99998 | 0.974509 | 0.150385 |
| 0.851696 | 0.409313 | 0.99998 | 0.851696 | 0.905661 | 0.053965 |
| 0.99998 | 0.99998 | 0.504103 | 0.99998 | 0.892165 | -0.107814 |
| 0.939903 | 0.99998 | 0.99998 | 0.939903 | 0.951843 | 0.011940 |
| 0.504103 | 0.99998 | 0.504103 | 0.495457 | 0.657915 | 0.153812 |
| 0.91464 | 0.409313 | 0.99998 | 0.91464 | 0.934886 | 0.020246 |
| 0.886851 | 0.99998 | 0.99998 | 0.886851 | 0.927211 | 0.040360 |
| 0.99998 | 0.99998 | 0.543698 | 0.99998 | 0.899157 | -0.100822 |
| 0.824377 | 0.99998 | 0.99998 | 0.668813 | 0.825976 | 0.001599 |
| 0.99998 | 0.409313 | 0.99998 | 0.99998 | 0.974509 | -0.025470 |
| 0.99998 | 0.409313 | 0.99998 | 0.99998 | 0.974509 | -0.025470 |
| 0.99998 | 0.409313 | 0.99998 | 0.99998 | 0.974509 | -0.025470 |
| 0.99998 | 0.99998 | 0.99998 | 0.99998 | 0.979737 | -0.020242 |
| 0.714645 | 0.99998 | 0.99998 | 0.668813 | 0.825976 | 0.111331 |
| 0.738655 | 0.824124 | 0.738655 | 0.738655 | 0.810697 | 0.072042 |
| 0.99998 | 0.409313 | 0.99998 | 0.409313 | 0.700262 | -0.299717 |
| 0.543698 | 0.409313 | 0.504103 | 0.543698 | 0.675085 | 0.131387 |
| 0.848436 | 0.848436 | 0.99998 | 0.848436 | 0.908033 | 0.059597 |
| 0.495457 | 0.99998 | 0.504103 | 0.495457 | 0.657915 | 0.162458 |
| 0.99998 | 0.409313 | 0.99998 | 0.409313 | 0.700262 | -0.299717 |
| 0.915084 | 0.668813 | 0.99998 | 0.915084 | 0.937388 | 0.022304 |
| 0.99998 | 0.409313 | 0.99998 | 0.999980 | 0.974509 | -0.025470 |
| 0.99998 | 0.409313 | 0.99998 | 0.999980 | 0.974509 | -0.025470 |
| 0.91922 | 0.99998 | 0.99998 | 0.919220 | 0.942240 | 0.023020 |
| 0.597255 | 0.99998 | 0.99998 | 0.597255 | 0.792751 | 0.195496 |
| 0.409313 | 0.409313 | 0.99998 | 0.409313 | 0.700262 | 0.290949 |
| 0.99998 | 0.99998 | 0.99998 | 0.668813 | 0.825976 | -0.174003 |
| 0.668813 | 0.409313 | 0.99998 | 0.668813 | 0.820748 | 0.151935 |
| 0.99998 | 0.409313 | 0.99998 | 0.999980 | 0.974509 | -0.025470 |
| 0.99998 | 0.409313 | 0.99998 | 0.999980 | 0.974509 | -0.025470 |
| 0.908604 | 0.99998 | 0.543698 | 0.495457 | 0.664907 | -0.243696 |
| 0.868421 | 0.409313 | 0.99998 | 0.868421 | 0.913426 | 0.045005 |
| 0.99998 | 0.99998 | 0.99998 | 0.999980 | 0.979737 | -0.020242 |

**Table 5.** Authentic Germplasm Composition and Part-specific Germplasm Prediction Results with HybridInception_ViT Pipeline.

| **True**<br>**value** | **LIMB_Pre** | **HEAD_Pre** | **TOTAL_Pre** | **Predict**<br>**Value** | **Deviation** |
| --- | --- | --- | --- | --- | --- |
| 0.824124 | 0.738655 | 0.824124 | 0.99998 | 0.892307 | 0.068183 |
| 0.851696 | 0.504103 | 0.99998 | 0.851696 | 0.863103 | 0.011407 |
| 0.99998 | 0.99998 | 0.824377 | 0.99998 | 0.920202 | -0.079777 |
| 0.939903 | 0.738655 | 0.738655 | 0.939903 | 0.834826 | -0.105076 |
| 0.504103 | 0.543698 | 0.99998 | 0.504103 | 0.710691 | 0.206588 |
| 0.91464 | 0.99998 | 0.824124 | 0.99998 | 0.920112 | 0.005472 |
| 0.886851 | 0.99998 | 0.99998 | 0.886851 | 0.931705 | 0.044854 |
| 0.99998 | 0.99998 | 0.824124 | 0.99998 | 0.920112 | -0.079867 |
| 0.824377 | 0.99998 | 0.504103 | 0.99998 | 0.806248 | -0.018128 |
| 0.99998 | 0.99998 | 0.99998 | 0.99998 | 0.982681 | -0.017298 |
| 0.99998 | 0.409313 | 0.99998 | 0.99998 | 0.919834 | -0.080145 |
| 0.99998 | 0.99998 | 0.824124 | 0.886851 | 0.869136 | -0.130843 |
| 0.99998 | 0.99998 | 0.824124 | 0.99998 | 0.920112 | -0.079867 |
| 0.714645 | 0.99998 | 0.99998 | 0.668813 | 0.833457 | 0.118812 |
| 0.738655 | 0.738655 | 0.738655 | 0.99998 | 0.861897 | 0.123242 |
| 0.99998 | 0.99998 | 0.99998 | 0.99998 | 0.982681 | -0.017298 |
| 0.543698 | 0.668813 | 0.738655 | 0.504103 | 0.631023 | 0.087325 |
| 0.848436 | 0.886851 | 0.824124 | 0.886851 | 0.857099 | 0.008663 |
| 0.495457 | 0.738655 | 0.738655 | 0.495457 | 0.634559 | 0.139102 |
| 0.99998 | 0.409313 | 0.99998 | 0.915084 | 0.881580 | -0.118399 |
| 0.915084 | 0.99998 | 0.738655 | 0.915084 | 0.851448 | -0.063635 |
| 0.99998 | 0.668813 | 0.738655 | 0.99998 | 0.854466 | -0.145513 |
| 0.99998 | 0.543698 | 0.824124 | 0.99998 | 0.871563 | -0.128416 |
| 0.91922 | 0.543698 | 0.939903 | 0.99998 | 0.912757 | -0.006462 |
| 0.597255 | 0.738655 | 0.738655 | 0.886851 | 0.810921 | 0.213666 |
| 0.409313 | 0.99998 | 0.824124 | 0.99998 | 0.920112 | 0.510799 |
| 0.99998 | 0.99998 | 0.99998 | 0.99998 | 0.982681 | -0.017298 |
| 0.668813 | 0.99998 | 0.824124 | 0.99998 | 0.920112 | 0.251299 |
| 0.99998 | 0.668813 | 0.99998 | 0.99998 | 0.947445 | -0.052534 |
| 0.99998 | 0.91464 | 0.824124 | 0.99998 | 0.911032 | -0.088947 |
| 0.908604 | 0.99998 | 0.824124 | 0.495457 | 0.692774 | -0.215829 |
| 0.868421 | 0.668813 | 0.99998 | 0.99998 | 0.947445 | 0.079024 |
| 0.99998 | 0.99998 | 0.99998 | 0.99998 | 0.982681 | -0.017298 |

For the InceptionV3-based pipeline, the optimal fitting equation was:

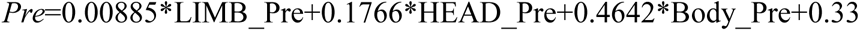

Fitting results showed moderate but meaningful consistency with the genomic germplasm composition:

1. Proportion of predictions with absolute error <10%: 57.57%
2. Proportion of predictions with absolute error<15%: 69.69%
3. Accuracy for individuals with both predicted and true germplasm purit<u>y</u> >90%: 68.42%

For the HybridInceptionViT pipeline, the fitted equation was:

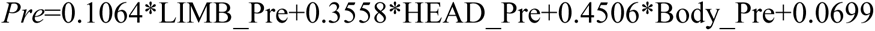

The hybrid model achieved noticeably improved fitting performance:

1. Proportion of predictions with absolute error <10%: 63.63%
2. Proportion of predictions with absolute error <15%: 87.87%
3. Accuracy for individuals with both predicted and true germplasm purit<u>y</u> >90%: 63.15%

### 3.7. Independent external testing of the germplasm identification pipeline

To further verify the reliability and generalization ability, the constructed pipeline was externally tested using an independent test dataset of eight individuals. For each individual, five images were randomly selected for prediction, and the result with the highest confidence was taken as the final phenotypic estimate. Germplasm composition was then inferred using the pre-established fitting equation.

Testing results based on the Inception-V3 model **(Table 6)**:

1. Proportion with absolute error <10%: 50%
2. Proportion with absolute error <15%: 75%
3. Accuracy for individuals with both predicted and true germplasm purity >90%: 20%

**Table 6.**
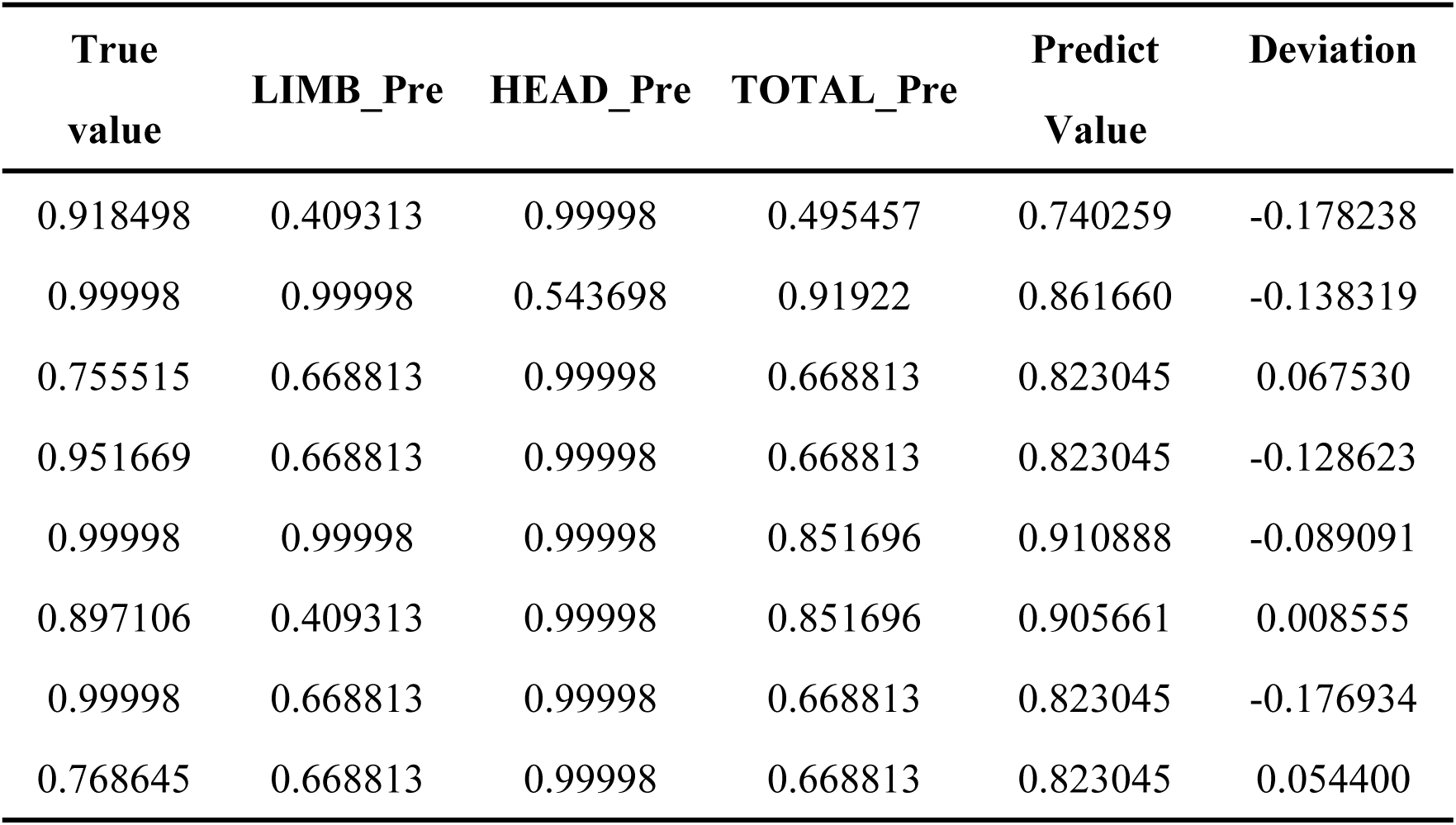
Validation Result with Inception_V3 Pipeline.

| True<br>value | LIMB_Pre | HEAD_Pre | TOTAL_Pre | Predict<br>Value | Deviation |
| --- | --- | --- | --- | --- | --- |
| 0.918498 | 0.409313 | 0.99998 | 0.495457 | 0.740259 | -0.178238 |
| 0.99998 | 0.99998 | 0.543698 | 0.91922 | 0.861660 | -0.138319 |
| 0.755515 | 0.668813 | 0.99998 | 0.668813 | 0.823045 | 0.067530 |
| 0.951669 | 0.668813 | 0.99998 | 0.668813 | 0.823045 | -0.128623 |
| 0.99998 | 0.99998 | 0.99998 | 0.851696 | 0.910888 | -0.089091 |
| 0.897106 | 0.409313 | 0.99998 | 0.851696 | 0.905661 | 0.008555 |
| 0.99998 | 0.668813 | 0.99998 | 0.668813 | 0.823045 | -0.176934 |
| 0.768645 | 0.668813 | 0.99998 | 0.668813 | 0.823045 | 0.054400 |

Testing results based on the HybridInceptionViT model **(Table 7)**:

1. Proportion with absolute error <10%: 75%
2. Proportion with absolute error <15%: 75%
3. Accuracy for individuals with both predicted and true germplasm purity >90%: 60%

**Table 7.** Validation Result with HybridInception_ViT Pipeline.

| True<br>value | LIMB_Pre | HEAD_Pre | TOTAL_Pre | Predict<br>Value | Deviation |
| --- | --- | --- | --- | --- | --- |
| 0.918498 | 0.668813 | 0.99998 | 0.99998 | 0.947445 | 0.028947 |
| 0.99998 | 0.908604 | 0.824124 | 0.848436 | 0.842104 | -0.157875 |
| 0.755515 | 0.99998 | 0.738655 | 0.495457 | 0.662364 | -0.093150 |
| 0.951669 | 0.543698 | 0.99998 | 0.99998 | 0.934133 | -0.017535 |
| 0.99998 | 0.886851 | 0.939903 | 0.668813 | 0.800045 | -0.199934 |
| 0.897106 | 0.99998 | 0.99998 | 0.99998 | 0.982681 | 0.085575 |
| 0.99998 | 0.99998 | 0.939903 | 0.886851 | 0.910330 | -0.089649 |
| 0.768645 | 0.543698 | 0.824124 | 0.851696 | 0.804747 | 0.036102 |

These outcomes were consistent with the internal fitting performance. The HybridInception ViT model demonstrated favorable and stable prediction, despite slight variation caused by the limited size of the external testing set.

## 4. Discussion

### 4.1. Integrated AI framework enables phenotype-genotype association in gayal

In this study, we developed an integrated artificial intelligence (AI) framework that couples image-based phenotypic recognition with population genomic analysis to quantify genomic admixture and germplasm composition in gayal. By constructing a curated dataset consisting of 6,245 morphological photographs and matched genomic sequences from 41 individuals, we established a standardized resource for systematic exploration of phenotype–genotype correlation in this endangered semi-feral bovine species. More importantly, our framework demonstrates the feasibility of inferring individual germplasm composition directly from visual phenotypes, representing a novel paradigm for livestock germplasm assessment.

Our results revealed abundant biologically meaningful genetic information embedded in animal morphological images. The high individual recognition accuracy and favorable germplasm fitting performance demonstrated by multiple models confirmed that visual traits such as body contour, head shape, limb morphology and coat characteristics contain sufficient discriminative genetic signals to distinguish conspecific individuals and infer germplasm composition.

Such morphological discrimination experiences have long been empirically used by local breeders and herdsmen for livestock identification and production, yet lacked systematic quantitative and standardized analytical methods (FAO 2012). Deep learning-based image recognition transformed these traditional qualitative morphological judgment into digital quantitative phenotypic traits, realizing precise digitization of morphological diversity. These digital phenotypes can comprehensively reflect genetic variation, individual development status and phenotypic differention patterns of livestock populations (Houle *et al*. 2010).

The anatomic part-based multi-modal pipeline constructed in this study fully exploited phenotypic differences among head, body, and limb regions, and improved the accuracy and stability of germplasm composition prediction through multi-dimensional feature fusion. Therefore, AI-assisted phenotypic mining technology endows morphological features as reliable biomarkers for individual identification and germplasm evaluation.

For rare and endangered livestock such as gayal, large-scale genomic screening is frequently constrained by high sequencing cost, complex welfare procedures, and field sampling in harsh geographical environments like remote mountainous regions where animals are often maintained under semi-free-ranging conditions (FAO 2015). Compared with traditional genome sequencing-dependent germplasm identification approaches, this image-based AI strategy offers a rapid, non-invasive, and easy-to-implement solution for preliminary germplasm evaluation. These advantages make it particularly suitable for large-scale on-site germplasm screening, providing convenient technical support for efficiency of conservation-oriented breeding programs (Distante *et al*. 2025). In addition, image-based preliminary screening can prioritize individuals with distinctive genetic backgrounds for subsequent in-depth genomic investigation, greatly reducing unnecessary sequencing expenditure and optimizing conservation resource allocation.

### 4.2. HybridInceptionViT improves phenotypic feature learning from morphological images

The core methodological innovation of this study lies in the construction of the HybridInceptionViT model, which synergistically integrates CNN convolutional architectures and transformer self-attention mechanisms to improve the extraction of morphological features from animal body images.

CNN models are highly effective at capturing local visual features such as edges, textures, and fine structural details, which are critical for characterizing animal morphological traits (Lecun *et al*. 1998; Krizhevsky *et al*. 2012). However, CNNs are relatively weak in modeling long-distance spatial dependencies and global holistic morphological relationships within images (Dosovitskiy *et al*. 2020). In contrast, ViT architectures excel at mining global image contextual information through self-attention mechanisms on ultra-large training datasets (Dosovitskiy *et al*. 2020).

Accordingly, we innovatively constructed the HybridInceptionViT model. Through parallel feature extraction and effective fusion of Inception multi-scale convolution module and ViT transformer module, it achieved simultaneous high-precision extraction of local fine features and global structural features. The Inception branch strengthened the recognition of delicate local phenotypic differences of gayal individuals, while the ViT branch mined overall body contour and global morphological association through the self-attention mechanism (Szegedy *et al*. 2015; Dosovitskiy *et al*. 2020). The complementary fusion of dual feature representations significantly improved its ability to interpret complex biological morphological variations under natural field conditions, parallel to the combination of Resnet50 and ViT (Peng *et al*. 2021). Furthermore, this hybrid architecture adopted wll to animal phenotypic datasets with limited sample sizes, a distinguished advantage observed in other hybrid model (Touvron *et al*. 2021a). Collectively, the substantial performance improvement of HybridInceptionViT observed in this study suggests that CNN-Transformer hybrid architectures may become a promising direction for future AI-assisted applications in biological image analysis.

### 4.3. Practical implications for conservation and management of endangered livestock

Gathering phenomic data is relatively expensive and time consuming (Houle *et al*. 2010). From a conservation perspective, the proposed AI phenotype-genotype joint analysis framework can effectively solve help address several challenges associated with the management of endangered livestock. Combined with wild monitoring cameras and mobile photographing systems, automated image recognition can continuously track individual genetic composition, assisting scientific breeding planning to maintain population genetic diversity while improving desirable excellent traits .

More importantly, the methodological framework developed here has broad popularization applicability, and can be extended to various species particularly those with conservation concern. Most endangered livestock and wild ungulates suffer common conservation challenges, including limited population size, incomplete pedigree records, and difficulty in large-scale genomic sampling (Frankham *et al*. 2002; FAO 2010). Integrating computer vision with genomic analysis offers a universal low-cost strategy for addressing these limitations. Such integrated approaches will play an increasingly important role in biodiversity conservation, livestock germplasm resource protectyion, and sustainable utilization.

### 4.5. Limitations and future perspectives

Several limitations should be noted. First, the current dataset is restricted in Chinese population, which may limit the representativeness of phenotypic and genetic variations across wild and scattered gayal populations. Second, phenotypic traits are influenced by confounding factors such as age, sex, growth stage, nutritional level, and feeding environment, which were not explicitly modeled in the present model. Third, the correlation between image predicted values and genomic admixture proportions was fitted using linear regression, whereas the underlying phenotype-genotype association may follow more complex nonlinear regularoty patterns.

Future studies may address these limitations by expanding multi-population sample coverage, incorporating additional environmental and physiological covariates, and exploring advanced nonlinear multimodal algorithms to directly fuse image phenotypic features and genomic variation data. Such developments could further elevate the accuracy of phenotype-based germplasm prediction and systematically reveal the genetic mechanism regulating morphological variation.

## 5. Conclusion

This study demonstrates the feasibility of integrating deep learning computer vision and population genomic analysis for germplasm purity evaluation in an endangered semi-feral livestock species. By establishing a phenotype–genotype association framework and innovatively developing the HybridInceptionViT architecture, we provided a convenient, efficient and non-invasive approach for rapid on-site gayal germplasm screening. More broadly, this interdisciplinary AI-genomic strategy highlighted the potential of AI-assisted phenotyping to support conservation genetics and sustainable management of animal genetic resources.

## Acknowledgements

This work was supported by the National Key R&D Program of China (2021YFD1200904); the National Natural Science Foundation of China (32470654&31860305); and Special funds for central guidance of local scientific and technological development (202407AA110003). Y.L. was supported by the Young Academic and Technical Leader Raising Foundation of Yunnan Province (2018HB033) and the Young Top Talents of the Ten Thousand Talents Plan in Yunnan Province (YNWR-QNBJ-2018-124). Yunnan University Postgraduate Research and Innovation Fund (ZC-242410931). Yunnan University Postgraduate Research and Innovation Fund (KC-242410789).

## Ethical approval

The ear tissue collection was performed in strict accordance with the protocol approved by the Institutional Animal Care and Use Committee of Yunnan University (Approval No. YNU20251450).

---

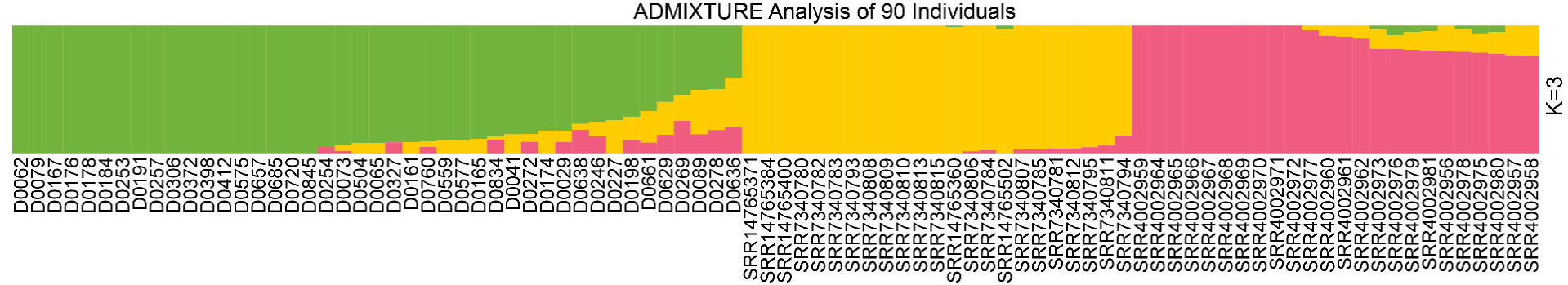

