## Supplemental information for "Bridging Morphology and Genomics: A rapid image-based assessment of genomic admixture in the endangered gayal (*Bos frontalis*)"

#### Appendix

##### Appendix A Species and Number of Individuals for Genomic ADMIXTURE Analysis

| Species | Number of Individuals |
|---|---|
| *Bos taurus* | 23 |
| *Bos indicus* | 24 |
| *Bos frontalis* | 67 |

**Appendix B Visualization Data of Q-matrix from ADMIXTURE Analysis**

| Name | Species | Germplasm Proportion [*Bos frontalis,Bos taurus,Bos indicus*] |
|---|---|---|
| SRR14765360 | *Bos taurus* | 0.009830 0.990160 0.000010 |
| SRR14765371 | *Bos taurus* | 0.000010 0.999980 0.000010 |
| SRR14765384 | *Bos taurus* | 0.000010 0.999980 0.000010 |
| SRR14765400 | *Bos taurus* | 0.000010 0.999980 0.000010 |
| SRR14765502 | *Bos taurus* | 0.027746 0.972244 0.000010 |
| SRR7340780 | *Bos taurus* | 0.000010 0.999980 0.000010 |
| SRR7340781 | *Bos taurus* | 0.000010 0.963220 0.036770 |
| SRR7340782 | *Bos taurus* | 0.000010 0.999980 0.000010 |
| SRR7340783 | *Bos taurus* | 0.000010 0.999980 0.000010 |
| SRR7340784 | *Bos taurus* | 0.000010 0.977590 0.022400 |
| SRR7340785 | *Bos taurus* | 0.000010 0.968990 0.031000 |
| SRR7340793 | *Bos taurus* | 0.000010 0.999980 0.000010 |
| SRR7340794 | *Bos taurus* | 0.000010 0.863957 0.136033 |
| SRR7340795 | *Bos taurus* | 0.000010 0.953800 0.046190 |
| SRR7340806 | *Bos taurus* | 0.000010 0.985558 0.014432 |
| SRR7340807 | *Bos taurus* | 0.000010 0.970222 0.029768 |
| SRR7340808 | *Bos taurus* | 0.000010 0.999980 0.000010 |
| SRR7340809 | *Bos taurus* | 0.000010 0.999980 0.000010 |
| SRR7340810 | *Bos taurus* | 0.000010 0.999980 0.000010 |
| SRR7340811 | *Bos taurus* | 0.009830 0.990160 0.000010 |
| SRR7340812 | *Bos taurus* | 0.000010 0.999980 0.000010 |
| SRR7340813 | *Bos taurus* | 0.000010 0.999980 0.000010 |
| SRR7340815 | *Bos taurus* | 0.000010 0.999980 0.000010 |
| SRR4002956 | *Bos indicus* | 0.000010 0.201352 0.798638 |
| SRR4002957 | *Bos indicus* | 0.000010 0.236026 0.763964 |
| SRR4002958 | *Bos indicus* | 0.000010 0.237209 0.762781 |

| | | | | |
|---|---|---|---|---|
| SRR4002959 | *Bos indicus* | 0.000010 | 0.000010 | 0.999980 |
| SRR4002960 | *Bos indicus* | 0.000010 | 0.079686 | 0.920304 |
| SRR4002961 | *Bos indicus* | 0.000010 | 0.085556 | 0.914434 |
| SRR4002962 | *Bos indicus* | 0.000010 | 0.102140 | 0.897850 |
| SRR4002964 | *Bos indicus* | 0.000010 | 0.000010 | 0.999980 |
| SRR4002965 | *Bos indicus* | 0.000010 | 0.000010 | 0.999980 |
| SRR4002966 | *Bos indicus* | 0.000010 | 0.000010 | 0.999980 |
| SRR4002967 | *Bos indicus* | 0.000010 | 0.000010 | 0.999980 |
| SRR4002968 | *Bos indicus* | 0.000010 | 0.000010 | 0.999980 |
| SRR4002969 | *Bos indicus* | 0.000010 | 0.000010 | 0.999980 |
| SRR4002970 | *Bos indicus* | 0.000010 | 0.000010 | 0.999980 |
| SRR4002971 | *Bos indicus* | 0.000010 | 0.000010 | 0.999980 |
| SRR4002972 | *Bos indicus* | 0.000010 | 0.000010 | 0.999980 |
| SRR4002973 | *Bos indicus* | 0.031891 | 0.149314 | 0.818796 |
| SRR4002975 | *Bos indicus* | 0.066109 | 0.146028 | 0.787862 |
| SRR4002976 | *Bos indicus* | 0.073568 | 0.110355 | 0.816077 |
| SRR4002977 | *Bos indicus* | 0.000010 | 0.035093 | 0.964897 |
| SRR4002978 | *Bos indicus* | 0.024342 | 0.181005 | 0.794653 |
| SRR4002979 | *Bos indicus* | 0.050412 | 0.140283 | 0.809305 |
| SRR4002980 | *Bos indicus* | 0.050768 | 0.170691 | 0.778541 |
| SRR4002981 | *Bos indicus* | 0.042545 | 0.154494 | 0.802960 |
| D0029 | *Bos frontalis* | 0.000010 | 0.940403 | 0.059587 |
| D0041 | *Bos frontalis* | 0.000010 | 0.962723 | 0.037267 |
| D0062 | *Bos frontalis* | 0.000010 | 0.999980 | 0.000010 |
| D0065 | *Bos frontalis* | 0.000010 | 0.999980 | 0.000010 |
| D0073 | *Bos frontalis* | 0.824124 | 0.087815 | 0.088061 |
| D0079 | *Bos frontalis* | 0.851696 | 0.148294 | 0.000010 |
| D0089 | *Bos frontalis* | 0.999980 | 0.000010 | 0.000010 |
| D0161 | *Bos frontalis* | 0.918498 | 0.080272 | 0.001229 |

| | | | | |
|---|---|---|---|---|
| D0165 | *Bos frontalis* | 0.939903 | 0.040942 | 0.019155 |
| D0167 | *Bos frontalis* | 0.999980 | 0.000010 | 0.000010 |
| D0174 | *Bos frontalis* | 0.504103 | 0.347839 | 0.148058 |
| D0176 | *Bos frontalis* | 0.914640 | 0.085350 | 0.000010 |
| D0178 | *Bos frontalis* | 0.886851 | 0.113139 | 0.000010 |
| D0184 | *Bos frontalis* | 0.999980 | 0.000010 | 0.000010 |
| D0198 | *Bos frontalis* | 0.824377 | 0.175613 | 0.000010 |
| D0227 | *Bos frontalis* | 0.999980 | 0.000010 | 0.000010 |
| D0246 | *Bos frontalis* | 0.999980 | 0.000010 | 0.000010 |
| D0253 | *Bos frontalis* | 0.999980 | 0.000010 | 0.000010 |
| D0269 | *Bos frontalis* | 0.714645 | 0.184639 | 0.100715 |
| D0272 | *Bos frontalis* | 0.738655 | 0.261335 | 0.000010 |
| D0278 | *Bos frontalis* | 0.755515 | 0.113090 | 0.131395 |
| D191 | *Bos frontalis* | 0.999980 | 0.000010 | 0.000010 |
| D254 | *Bos frontalis* | 0.543698 | 0.201710 | 0.254592 |
| D257 | *Bos frontalis* | 0.848436 | 0.061689 | 0.089875 |
| D306 | *Bos frontalis* | 0.495457 | 0.323529 | 0.181014 |
| D327 | *Bos frontalis* | 0.999980 | 0.000010 | 0.000010 |
| D372 | *Bos frontalis* | 0.951669 | 0.000010 | 0.048321 |
| D398 | *Bos frontalis* | 0.999980 | 0.000010 | 0.000010 |
| D412 | *Bos frontalis* | 0.999980 | 0.000010 | 0.000010 |
| D504 | *Bos frontalis* | 0.915084 | 0.000010 | 0.084906 |
| D559 | *Bos frontalis* | 0.999980 | 0.000010 | 0.000010 |
| D575 | *Bos frontalis* | 0.999980 | 0.000010 | 0.000010 |
| D577 | *Bos frontalis* | 0.999980 | 0.000010 | 0.000010 |
| D629 | *Bos frontalis* | 0.919220 | 0.080770 | 0.000010 |
| D636 | *Bos frontalis* | 0.897106 | 0.102884 | 0.000010 |
| D638 | *Bos frontalis* | 0.999980 | 0.000010 | 0.000010 |
| D657 | *Bos frontalis* | 0.896939 | 0.103051 | 0.000010 |

| | | | | |
|---|---|---|---|---|
| D661 | *Bos frontalis* | 0.597255 | 0.256507 | 0.146238 |
| D685 | *Bos frontalis* | 0.409313 | 0.388062 | 0.202625 |
| D720 | *Bos frontalis* | 0.768645 | 0.047947 | 0.183407 |
| D760 | *Bos frontalis* | 0.999980 | 0.000010 | 0.000010 |
| D834 | *Bos frontalis* | 0.668813 | 0.247548 | 0.083639 |
| D845 | *Bos frontalis* | 0.999980 | 0.000010 | 0.000010 |
| D0244 | *Bos frontalis* | 0.510282 | 0.477460 | 0.012258 |
| D0247 | *Bos frontalis* | 0.510253 | 0.477485 | 0.012262 |
| D0040 | *Bos frontalis* | 0.995168 | 0.004822 | 0.000010 |
| D0687 | *Bos frontalis* | 0.773155 | 0.226835 | 0.000010 |
| GS01 | *Bos frontalis* | 0.977521 | 0.022469 | 0.000010 |
| GS02 | *Bos frontalis* | 0.998738 | 0.000010 | 0.001252 |
| GS03 | *Bos frontalis* | 0.697414 | 0.201132 | 0.101454 |
| GS04 | *Bos frontalis* | 0.947810 | 0.052180 | 0.000010 |
| LS01 | *Bos frontalis* | 0.999980 | 0.000010 | 0.000010 |
| LS02 | *Bos frontalis* | 0.552645 | 0.424277 | 0.023079 |
| LS03 | *Bos frontalis* | 0.941162 | 0.058828 | 0.000010 |
| LS04 | *Bos frontalis* | 0.938914 | 0.061076 | 0.000010 |
| LS05 | *Bos frontalis* | 0.988727 | 0.011263 | 0.000010 |
| D882 | *Bos frontalis* | 0.999980 | 0.000010 | 0.000010 |
| D890 | *Bos frontalis* | 0.865929 | 0.072784 | 0.061287 |
| D2 | *Bos frontalis* | 0.763562 | 0.155960 | 0.080478 |
| D3 | *Bos frontalis* | 0.725330 | 0.181074 | 0.093596 |
| D5 | *Bos frontalis* | 0.658613 | 0.253738 | 0.087650 |
| D6 | *Bos frontalis* | 0.745410 | 0.162194 | 0.092396 |

**Appendix C Information of 114 individual genomic samples**

| Smple ID | Species | Source |
|---|---|---|
| SRR14765360 | Bos_taurus_taurus | CAN_Angus |
| SRR14765371 | Bos_taurus_taurus | CAN_Angus |
| SRR14765384 | Bos_taurus_taurus | CAN_Angus |
| SRR14765400 | Bos_taurus_taurus | CAN_Angus |
| SRR14765502 | Bos_taurus_taurus | CAN_Angus |
| SRR7340780 | Bos_taurus_taurus | CAN_Angus |
| SRR7340781 | Bos_taurus_taurus | CAN_Angus |
| SRR7340782 | Bos_taurus_taurus | CAN_Angus |
| SRR7340783 | Bos_taurus_taurus | CAN_Angus |
| SRR7340784 | Bos_taurus_taurus | CAN_Angus |
| SRR7340785 | Bos_taurus_taurus | CAN_Angus |
| SRR7340793 | Bos_taurus_taurus | CAN_Angus |
| SRR7340794 | Bos_taurus_taurus | CAN_Angus |
| SRR7340795 | Bos_taurus_taurus | CAN_Angus |
| SRR7340806 | Bos_taurus_taurus | CAN_Angus |
| SRR7340807 | Bos_taurus_taurus | CAN_Angus |
| SRR7340808 | Bos_taurus_taurus | CAN_Angus |
| SRR7340809 | Bos_taurus_taurus | CAN_Angus |
| SRR7340810 | Bos_taurus_taurus | CAN_Angus |
| SRR7340811 | Bos_taurus_taurus | CAN_Angus |
| SRR7340812 | Bos_taurus_taurus | CAN_Angus |
| SRR7340813 | Bos_taurus_taurus | CAN_Angus |
| SRR7340815 | Bos_taurus_taurus | CAN_Angus |
| SRR4002956 | Bos_taurus_indicus | U.S._Brahman |
| SRR4002957 | Bos_taurus_indicus | U.S._Brahman |
| SRR4002958 | Bos_taurus_indicus | U.S._Brahman |
| SRR4002959 | Bos_taurus_indicus | U.S._Brahman |

| | | |
|---|---|---|
| SRR4002960 | Bos_taurus_indicus | U.S._Brahman |
| SRR4002961 | Bos_taurus_indicus | U.S._Brahman |
| SRR4002962 | Bos_taurus_indicus | U.S._Brahman |
| SRR4002964 | Bos_taurus_indicus | U.S._Brahman |
| SRR4002965 | Bos_taurus_indicus | U.S._Brahman |
| SRR4002966 | Bos_taurus_indicus | U.S._Brahman |
| SRR4002967 | Bos_taurus_indicus | U.S._Brahman |
| SRR4002968 | Bos_taurus_indicus | U.S._Brahman |
| SRR4002969 | Bos_taurus_indicus | U.S._Brahman |
| SRR4002970 | Bos_taurus_indicus | U.S._Brahman |
| SRR4002971 | Bos_taurus_indicus | U.S._Brahman |
| SRR4002972 | Bos_taurus_indicus | U.S._Brahman |
| SRR4002973 | Bos_taurus_indicus | U.S._Brahman |
| SRR4002975 | Bos_taurus_indicus | U.S._Brahman |
| SRR4002976 | Bos_taurus_indicus | U.S._Brahman |
| SRR4002977 | Bos_taurus_indicus | U.S._Brahman |
| SRR4002978 | Bos_taurus_indicus | U.S._Brahman |
| SRR4002979 | Bos_taurus_indicus | U.S._Brahman |
| SRR4002980 | Bos_taurus_indicus | U.S._Brahman |
| SRR4002981 | Bos_taurus_indicus | U.S._Brahman |
| D0029 | Bos_frontalis | Nujiang_gayal |
| D0041 | Bos_frontalis | Nujiang_gayal |
| D0062 | Bos_frontalis | Nujiang_gayal |
| D0065 | Bos_frontalis | Nujiang_gayal |
| D0073 | Bos_frontalis | Nujiang_gayal |
| D0079 | Bos_frontalis | Nujiang_gayal |
| D0089 | Bos_frontalis | Nujiang_gayal |
| D0161 | Bos_frontalis | Nujiang_gayal |
| D0165 | Bos_frontalis | Nujiang_gayal |

| | | |
|---|---|---|
| D0167 | Bos_frontalis | Nujiang_gayal |
| D0174 | Bos_frontalis | Nujiang_gayal |
| D0176 | Bos_frontalis | Nujiang_gayal |
| D0178 | Bos_frontalis | Nujiang_gayal |
| D0184 | Bos_frontalis | Nujiang_gayal |
| D0198 | Bos_frontalis | Nujiang_gayal |
| D0227 | Bos_frontalis | Nujiang_gayal |
| D0246 | Bos_frontalis | Nujiang_gayal |
| D0253 | Bos_frontalis | Nujiang_gayal |
| D0269 | Bos_frontalis | Nujiang_gayal |
| D0272 | Bos_frontalis | Nujiang_gayal |
| D0278 | Bos_frontalis | Nujiang_gayal |
| D191 | Bos_frontalis | Nujiang_gayal |
| D254 | Bos_frontalis | Nujiang_gayal |
| D257 | Bos_frontalis | Nujiang_gayal |
| D306 | Bos_frontalis | Nujiang_gayal |
| D327 | Bos_frontalis | Nujiang_gayal |
| D372 | Bos_frontalis | Nujiang_gayal |
| D398 | Bos_frontalis | Nujiang_gayal |
| D412 | Bos_frontalis | Nujiang_gayal |
| D504 | Bos_frontalis | Nujiang_gayal |
| D559 | Bos_frontalis | Nujiang_gayal |
| D575 | Bos_frontalis | Nujiang_gayal |
| D577 | Bos_frontalis | Nujiang_gayal |
| D629 | Bos_frontalis | Nujiang_gayal |
| D636 | Bos_frontalis | Nujiang_gayal |
| D638 | Bos_frontalis | Nujiang_gayal |
| D657 | Bos_frontalis | Nujiang_gayal |
| D661 | Bos_frontalis | Nujiang_gayal |

| | | |
|---|---|---|
| D685 | Bos_frontalis | Nujiang_gayal |
| D720 | Bos_frontalis | Nujiang_gayal |
| D760 | Bos_frontalis | Nujiang_gayal |
| D834 | Bos_frontalis | Nujiang_gayal |
| D845 | Bos_frontalis | Nujiang_gayal |
| D0244 | Bos_frontalis | Nujiang_gayal |
| D0247 | Bos_frontalis | Nujiang_gayal |
| D0040 | Bos_frontalis | Nujiang_gayal |
| D0687 | Bos_frontalis | Nujiang_gayal |
| GS01 | Bos_frontalis | Nujiang_gayal |
| GS02 | Bos_frontalis | Nujiang_gayal |
| GS03 | Bos_frontalis | Nujiang_gayal |
| GS04 | Bos_frontalis | Nujiang_gayal |
| LS01 | Bos_frontalis | Nujiang_gayal |
| LS02 | Bos_frontalis | Nujiang_gayal |
| LS03 | Bos_frontalis | Nujiang_gayal |
| LS04 | Bos_frontalis | Nujiang_gayal |
| LS05 | Bos_frontalis | Nujiang_gayal |
| D882 | Bos_frontalis | Nujiang_gayal |
| D890 | Bos_frontalis | Nujiang_gayal |
| D2 | Bos_frontalis | Nujiang_gayal |
| D3 | Bos_frontalis | Nujiang_gayal |
| D5 | Bos_frontalis | Nujiang_gayal |
| D6 | Bos_frontalis | Nujiang_gayal |

**Appendix D Two Sampling Campaigns**

| Data Set | Number of Individuals | Number of Pictures |
|---|---|---|
| First | 15 | 715 |
| Second | 52 | 6245 |

带格式表格[马俊涛]

**Appendix E Genomic ADMIXTURE results at K=3 and K=4**

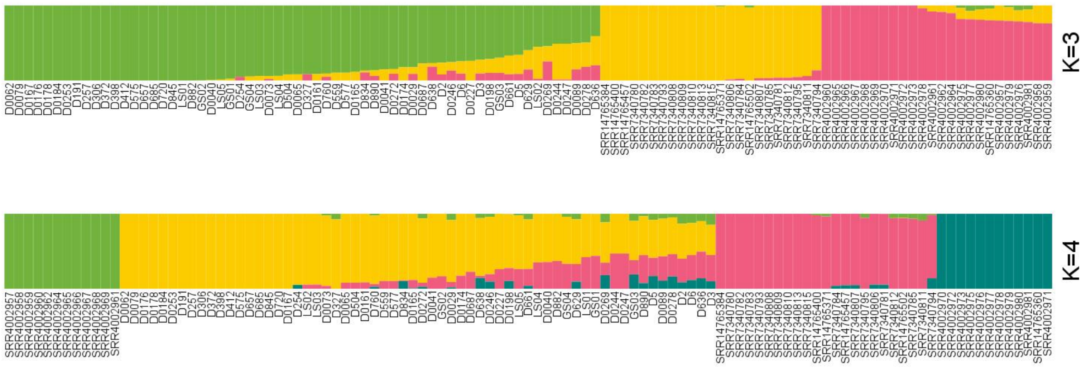


**Appendix F Image Data of 12 Individuals for Pre-training from the First Sampling of 15 Individuals**

| Name | Number of images in the training set | Number of images in the test set | Total number of images |
|---|---|---|---|
| D0033 | 44 | 12 | 56 |
| D0085 | 47 | 12 | 59 |
| D0088 | 38 | 11 | 49 |
| D0091 | 59 | 15 | 74 |
| D0189 | 38 | 9 | 47 |
| D0199 | 37 | 9 | 46 |
| D0227 | 24 | 6 | 30 |
| D0236 | 29 | 6 | 35 |
| D0446 | 59 | 14 | 73 |
| D0682 | 55 | 13 | 68 |
| D0733 | 87 | 21 | 108 |
| D0882 | 32 | 8 | 40 |
| D0174 | None | None | 3 |
| D0196 | None | None | 7 |
| D0299 | None | None | 20 |

Data from 3 individuals were excluded from the training set due to an insufficient number of images.

**Appendix G Number of Images Corresponding to 33 Individuals in the Head Training Set**

| Name | Number of images in the training set | Number of images in the test set | Total number of images |
|---|---|---|---|
| D0029 | 179 | 42 | 221 |
| D0041 | 28 | 7 | 35 |
| D0062 | 79 | 19 | 98 |
| D0073 | 99 | 24 | 123 |
| D0089 | 113 | 27 | 140 |
| D0161 | 30 | 8 | 38 |
| D0165 | 32 | 8 | 40 |
| D0167 | 61 | 15 | 76 |
| D0174 | 59 | 15 | 74 |
| D0176 | 65 | 15 | 80 |
| D0178 | 263 | 63 | 326 |
| D0184 | 144 | 32 | 176 |
| D0191 | 65 | 15 | 80 |
| D0198 | 69 | 17 | 86 |
| D0227 | 78 | 18 | 96 |
| D0253 | 78 | 18 | 96 |
| D0269 | 40 | 11 | 51 |
| D0272 | 66 | 16 | 82 |
| D0278 | 78 | 18 | 96 |
| D0306 | 78 | 18 | 96 |
| D0327 | 50 | 12 | 62 |
| D0372 | 34 | 8 | 42 |
| D0398 | 59 | 15 | 74 |

| | | | |
|---|---|---|---|
| D0504 | 57 | 14 | 71 |
| D0629 | 78 | 18 | 96 |
| D0636 | 58 | 14 | 72 |
| D0657 | 78 | 18 | 96 |
| D0661 | 27 | 6 | 33 |
| D0685 | 77 | 18 | 95 |
| D0720 | 66 | 16 | 82 |
| D0760 | 77 | 18 | 95 |
| D0834 | 47 | 12 | 59 |
| D0845 | 65 | 15 | 80 |

**Appendix H Number of Images Corresponding to 33 Individuals in the Body Training Set**

| Name | Number of images in the training set | Number of images in the test set | Total number of images |
|---|---|---|---|
| D0029 | 78 | 18 | 96 |
| D0041 | 27 | 8 | 35 |
| D0062 | 80 | 18 | 98 |
| D0073 | 99 | 24 | 123 |
| D0089 | 113 | 27 | 140 |
| D0161 | 35 | 9 | 44 |
| D0165 | 33 | 9 | 42 |
| D0167 | 61 | 15 | 76 |
| D0174 | 59 | 15 | 74 |
| D0176 | 65 | 15 | 80 |
| D0178 | 266 | 60 | 326 |
| D0184 | 143 | 33 | 176 |
| D0191 | 65 | 15 | 80 |
| D0198 | 69 | 17 | 86 |
| D0227 | 78 | 18 | 96 |
| D0253 | 78 | 18 | 96 |
| D0269 | 41 | 10 | 51 |
| D0272 | 66 | 16 | 82 |
| D0278 | 78 | 18 | 96 |
| D0306 | 78 | 18 | 96 |
| D0327 | 50 | 12 | 62 |
| D0372 | 33 | 9 | 42 |
| D0398 | 59 | 15 | 74 |
| D0504 | 59 | 15 | 74 |
| D0629 | 78 | 18 | 96 |

| | | | |
|---|---|---|---|
| D0636 | 58 | 14 | 72 |
| D0657 | 78 | 18 | 96 |
| D0661 | 27 | 6 | 33 |
| D0685 | 77 | 18 | 95 |
| D0720 | 66 | 16 | 82 |
| D0760 | 78 | 18 | 96 |
| D0834 | 47 | 12 | 59 |
| D0845 | 65 | 15 | 80 |

**Appendix I Number of Images Corresponding to 32 Individuals in the LIMB Training Set**

| Name | Number of images in the training set | Number of images in the test set | Total number of images |
|---|---|---|---|
| D0029 | 169 | 40 | 209 |
| D0062 | 78 | 20 | 98 |
| D0073 | 99 | 24 | 123 |
| D0089 | 113 | 27 | 140 |
| D0161 | 35 | 9 | 44 |
| D0165 | 34 | 8 | 42 |
| D0167 | 61 | 15 | 76 |
| D0174 | 59 | 15 | 74 |
| D0176 | 65 | 15 | 80 |
| D0178 | 263 | 63 | 326 |
| D0184 | 133 | 33 | 166 |
| D0191 | 61 | 15 | 76 |
| D0198 | 68 | 18 | 86 |
| D0227 | 63 | 15 | 78 |
| D0253 | 78 | 18 | 96 |
| D0269 | 41 | 10 | 51 |
| D0272 | 65 | 16 | 81 |
| D0278 | 78 | 18 | 96 |
| D0306 | 78 | 18 | 96 |
| D0327 | 50 | 12 | 62 |
| D0372 | 34 | 8 | 42 |
| D0398 | 59 | 15 | 74 |
| D0504 | 57 | 14 | 71 |
| D0629 | 78 | 18 | 96 |
| D0636 | 58 | 14 | 72 |
| D0657 | 78 | 18 | 96 |

| | | | |
|---|---|---|---|
| D0661 | 19 | 4 | 23 |
| D0685 | 77 | 18 | 95 |
| D0720 | 66 | 16 | 82 |
| D0760 | 78 | 18 | 96 |
| D0834 | 47 | 12 | 59 |
| D0845 | 58 | 15 | 73 |

**Appendix J Part-specific training sets**

| Training Set | Number of individuals | Number of pictures |
|---|---|---|
| Head | 33 | 3067 |
| Body | 33 | 2954 |
| Limb | 32 | 2929 |

The limb training set lacks one corresponding individual due to its recumbent posture.

**Appendix K Efficientnet Training Result**

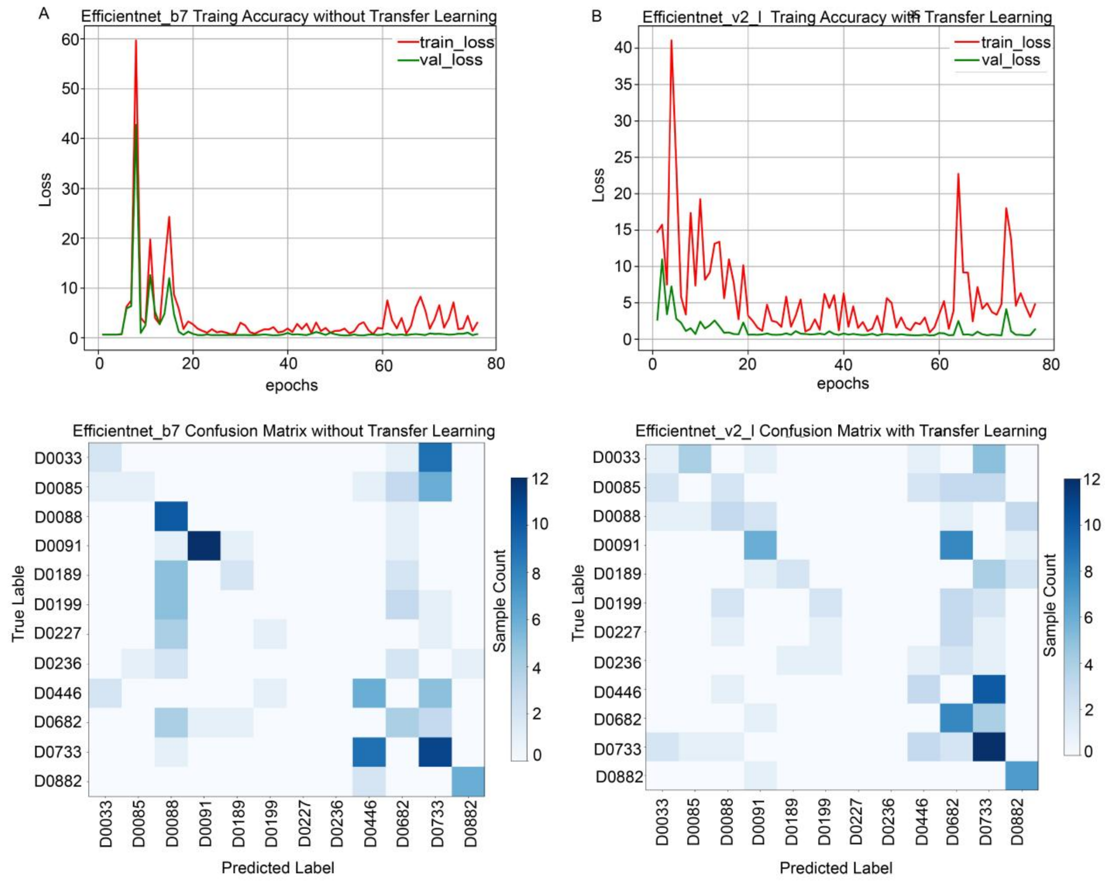

### Appendix L Grad-CAM visualization of body parts for individual D0029

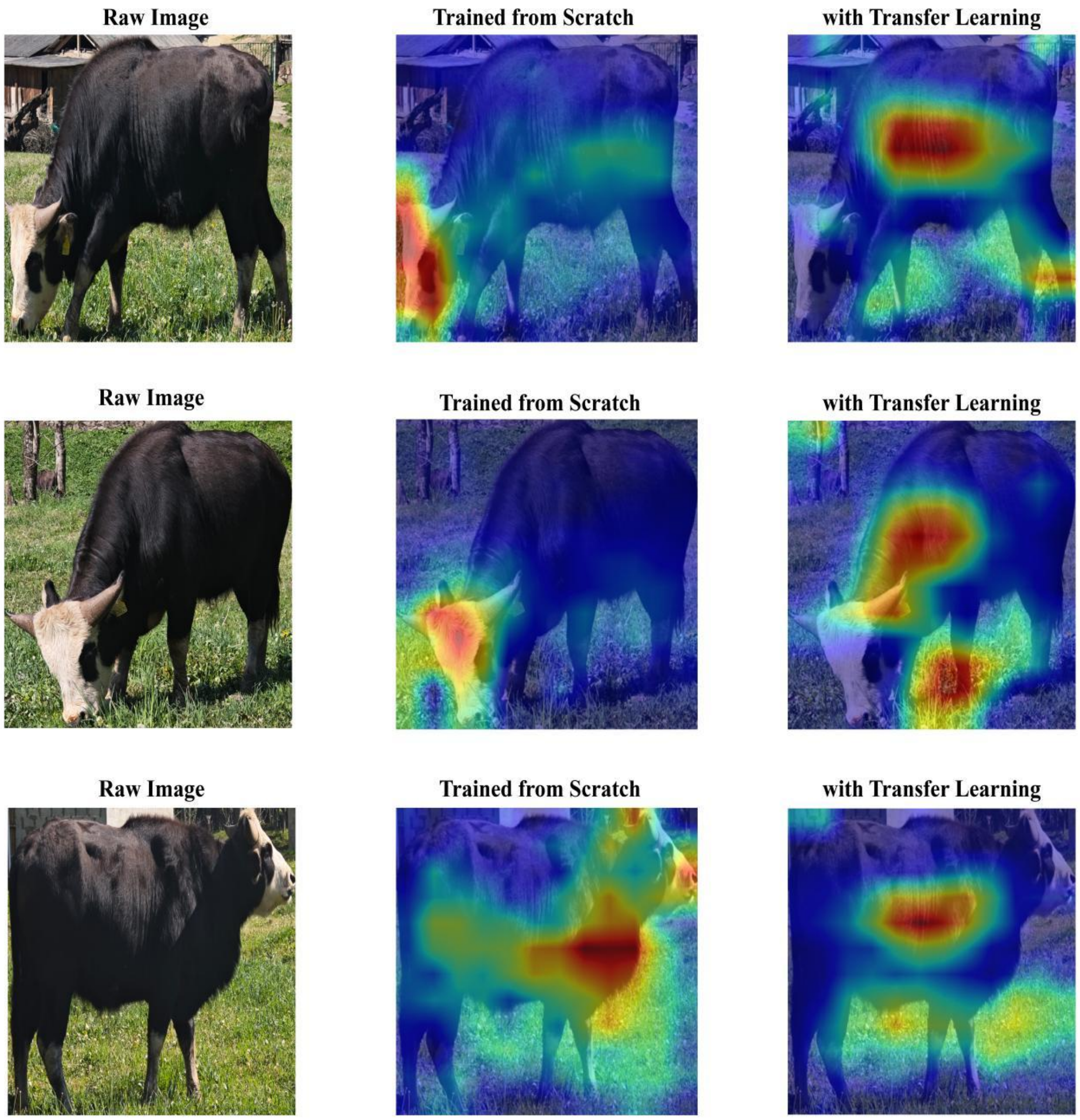

#### Appendix M Grad-CAM visualization of body parts for individual D0073

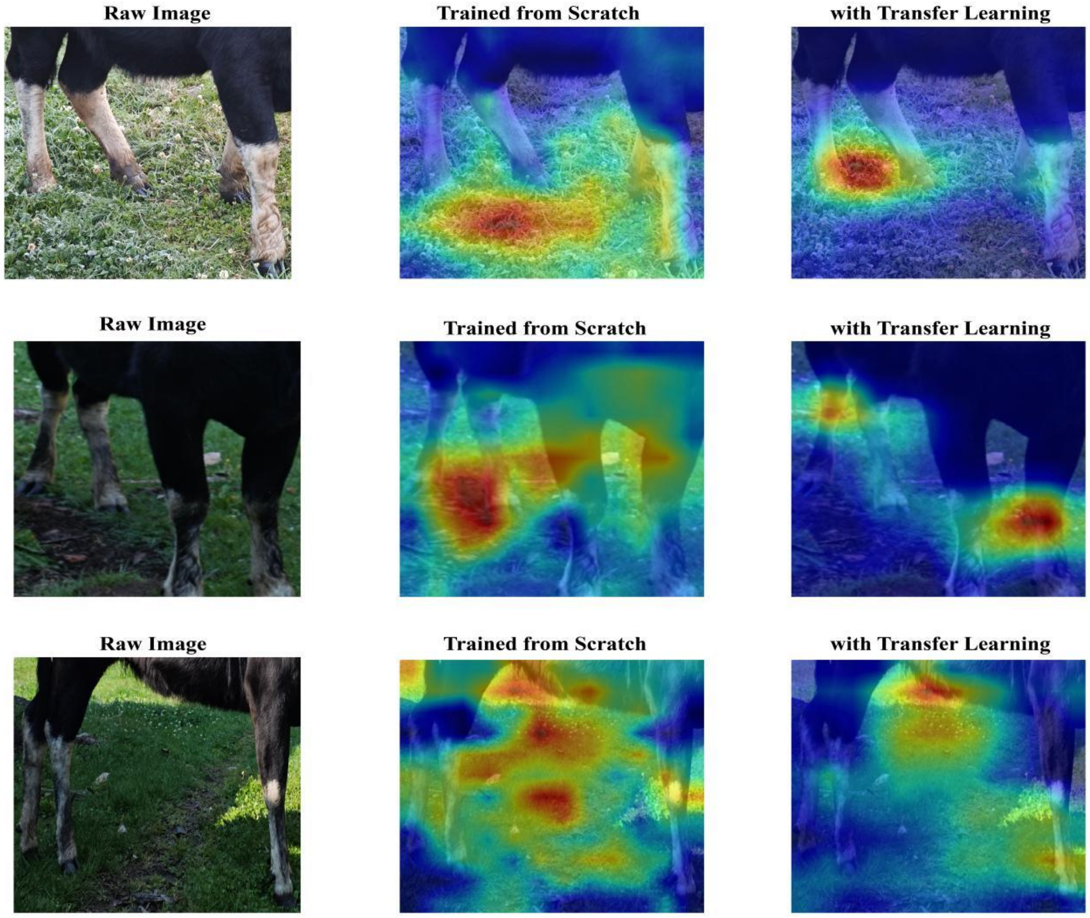
